# Mitochondria preserve an autarkic one-carbon cycle to confer growth-independent cancer cell migration and metastasis

**DOI:** 10.1101/2021.05.27.445928

**Authors:** Nicole Kiweler, Catherine Delbrouck, Vitaly I. Pozdeev, Laura Neises, Leticia Soriano-Baguet, Kim Eiden, Feng Xian, Mohaned Benzarti, Lara Haase, Eric Koncina, Maryse Schmoetten, Christian Jaeger, Muhammad Zaeem Noman, Alexei Vazquez, Bassam Janji, Gunnar Dittmar, Dirk Brenner, Elisabeth Letellier, Johannes Meiser

## Abstract

Progression of primary cancer to metastatic disease is the most common cause of death in cancer patients with minimal treatment options available. Canonical drugs target mainly the proliferative capacity of cancer cells, which often leaves slow-proliferating, persistent cancer cells unaffected. Metabolic determinants that contribute to growth-independent functions supporting resistance and metastatic dissemination are still poorly understood.

In the present study, we revealed that antifolate treatment results in an uncoupled and autarkic mitochondrial one-carbon (1C) metabolism allowing sustained serine catabolism and formate overflow when cytosolic 1C metabolism is impaired. Interestingly, antifolate dependent growth-arrest did not correlate with decreased migration capacity. Therefore, using the antifolate Methotrexate as a tool compound allowed us to disentangle proliferation and migration to profile the metabolic phenotype of migrating (growth-arrested) cells. Supported by an increased NAD/NADH ratio, we observed increased serine *de novo* synthesis and increased serine catabolism to formate.

Consequently, inhibition of serine *de novo* synthesis using the competitive PHGDH-inhibitor BI-4916 or direct inhibition of mitochondrial 1C metabolism reduced cancer cell migration. Using an orthotopic breast cancer model, we show that sole inhibition of mitochondrial serine catabolism does not affect primary tumor growth but strongly inhibits pulmonary metastasis.

We conclude that mitochondrial 1C metabolism, despite being dispensable for proliferative capacities, confers an advantage to cancer cells by supporting their motility potential.

Our results improve our understanding of 1C metabolism and of metabolic determinants that support the process of cancer cell migration and metastasis.

## Introduction

The survival rates of patients harboring primary tumors steadily increase due to targeted treatment schemes. However, in case of tumor relapse and metastatic disease, effective therapies are mostly lacking with the consequence that secondary tumors account for the majority of cancer deaths [1]. Classical chemotherapeutic approaches to counteract cancer growth aim to target biomass production by interfering with the synthesis of proteins, lipids, and nucleotides and thereby directly interfere with central pathways of cancer cell metabolism [2, 3]. Prospectively, therapeutic approaches targeting metabolic pathways that also support the invasive and migratory properties of cancer cells would help to prevent metastatic progression of the initial disease. However, profound knowledge of specific metabolic dependencies that support growth-independent processes during the metastatic cascade, are only starting to be understood [4].

Nucleotide synthesis is an essential requirement for cancer cell proliferation. Consequently, antifolates such as Methotrexate (MTX) and related compounds have been proven as successful chemotherapeutics for several decades. Although differences between the various antifolates exist, a shared mode of action is the inhibition of one-carbon (1C) metabolism which provides essential building blocks for thymidylate and purine synthesis. Additionally, 1C metabolism has implications in supporting methylation reactions, glutathione, heme, sphingolipid and protein synthesis [5]. 1C metabolism describes a metabolic cycle that is spread across mitochondrion and cytoplasm and it is mainly fueled by the non-essential amino acid serine. De-novo serine synthesis is catalyzed by the rate limiting enzyme phosphoglycerate dehydrogenase (PHGDH) which is amplified in different tumors [6-8]. Hence, targeting serine synthesis, serine catabolism or serine availability by starvation or inhibition of PHDGH has been proven successful to inhibit cancer progression [9-17]. To support the various anabolic programs, cytosolic 1C metabolism is of particular importance, while mitochondrial 1C metabolism is dispensable due to the reversibility of the cytosolic 1C metabolism [16]. In fact, depending on intracellular folate levels, cytosolic 1C metabolism can be the dominant route to provide one-carbon units for nucleotide synthesis [17]. In that light, reasons for conserved mitochondrial 1C metabolism are still not fully understood. Previous evidence suggests implications of mitochondrial 1C metabolism for mitochondrial translation [18, 19]. One additional explanation resides in the observation that mitochondrial derived formate lowers the pool of free cytosolic folates by increasing the 10-CHO-THF pool thereby protecting it from degradation. Upon MTX, degradation is facilitated as folates accumulate in the oxidized DHF form, impeding the 10-CHO-THF synthetase activity of MTHFD1 [20]. However, if mitochondrial folate-dependent 1C metabolism and formate production remains functional under conditions of cytosolic 1C metabolism impairment and whether mitochondrial 1C metabolism itself confers additional functions that contribute to tumorigenesis remains unknown.

We have previously shown that a reduction of biomass demand can result in increased formate release from cancer cells to their surrounding microenvironment [21] and that the rate of serine catabolism to formate is significantly increased in adenomas of the small intestine and breast tumors compared to normal tissue [22]. Importantly, enhanced formate release rates and increased extracellular formate concentrations have been shown to increase the invasiveness of glioblastoma cells *in vitro* [22].

Apart from our previous findings on formate overflow and its potential implications for cancer cell invasion [21, 22], other reports substantiate a role of serine metabolism in the context of metastasis [7, 14, 23-25]. However, such studies were so far focused on the relevance of serine *de novo* synthesis via PHGDH or extracellular serine availability for metastatic outgrowth (proliferative aspect) in the secondary tissue [7, 14, 23-25]. In contrast, whether mitochondrial serine catabolism also has a growth-independent role in promoting cancer cell motility that potentially increases the rate of cancer cell escape from the primary tumor has not been addressed so far. Being able to identify and target metabolic programs that promote the early steps of the metastatic cascade would help to contain the tumor, a clinical phenotype with more favorable therapeutic outcome compared to advanced metastatic stages.

In the present study, we observed that antifolates, albeit being strongly cytostatic, did not decrease the cell’s motility potential. Using MTX as a tool compound to selectively inhibit cytosolic 1C metabolism, we show that mitochondrial 1C metabolism runs as an autarkic, compartmentalized pathway that is important to drive cancer cell motility. In contrast, the cytosolic part of 1C metabolism, which is essential for nucleotide synthesis during proliferation, is dispensable for the cellular motility potential. Hence, we provide mechanistic evidence that one selective advantage of mitochondrial 1C metabolism is to support cell motility. We corroborate this hypothesis in an orthotopic breast cancer model. Here, we demonstrate that ablating mitochondrial 1C metabolism effectively blocks mitochondrial serine catabolism but does not affect primary tumor growth. Yet, it strongly inhibits pulmonary metastasis formation.

## Results

### Inhibition of Anabolic Synthesis Routes Differentially Impacts the Motility Potential of Cancer Cells

To assess the relative importance of different anabolic pathways for cancer cell migration, we employed a panel of metabolic perturbations targeting glycolysis, mitochondrial electron transport chain (ETC) as well as lipid, protein, and nucleotide synthesis (**Figure 1A**). As expected, the chosen metabolic interventions inhibited proliferation and affected cell cycle progression of MDA-MB-468 breast cancer cells with no to mild toxic effects after 48 h treatment (**Figure 1B-C, S1A**). Inhibition of glycolysis, ETC, lipid, and protein synthesis significantly reduced migration of MDA-MB-468 cells (**Figure 1D**). Surprisingly and in contrast to all other growth-inhibiting conditions, inhibition of nucleotide synthesis with various drugs did not diminish migration of MDA-MB-468 cells (**Figure 1E**). Calculation of the area under curve (AUC) allowed us to quantitatively compare wound closure over time (**Figure 1D, E**) and confirmed that wound closure was significantly reduced upon treatment with rotenone (Rot), galactose (Gal), simvastatin (SIM) and sirolimus (rapamycin, SIR). However, treatment with the antifolates methotrexate (MTX) and pemetrexed (PEM), as well as with hydroxyurea (HU) and clofarabine (CLO) did not affect or did even significantly increase wound closure in scratch assays in comparison to fully proliferative, untreated control cells. While cell migration correlates with reduced cell growth for all other metabolic perturbations, nucleotide synthesis inhibition does not reduce cell migration in correlation to growth inhibition (**Figure 1F, G**). Sustained migration in response to nucleotide synthesis inhibition was verified in LN229 glioblastoma and 4T1 breast cancer cells (**Figure S1B-D**). Of note, using dialyzed or normal FBS did not change the result (**Figure S1E**). This observation was also confirmed in trans-well migration assays using Boyden chambers (**Figure 1H, S1F, G**). Furthermore, using ECM-collagen coating in trans-well assay, we found that nucleotide synthesis inhibition also had no impact on the invasive capacity of MDA-MB-468 cells (**Figure 1H**), LN229 and 4T1 cells (**Figure S1F, G**). To validate if MTX resistance results in a pro-migratory effect, we generated MTX-resistant MDA-MB-468 cells by long-term cultivation in 50 nM MTX for 2 months. The resulting MTX-resistant MDA-MB-468 cells proliferate in the presence of increasing concentrations of MTX (**Figure 1I**) and display enhanced migratory capacity compared to the parental MDA-MB-468 cell line (**Figure 1J**).

**Figure 1:**
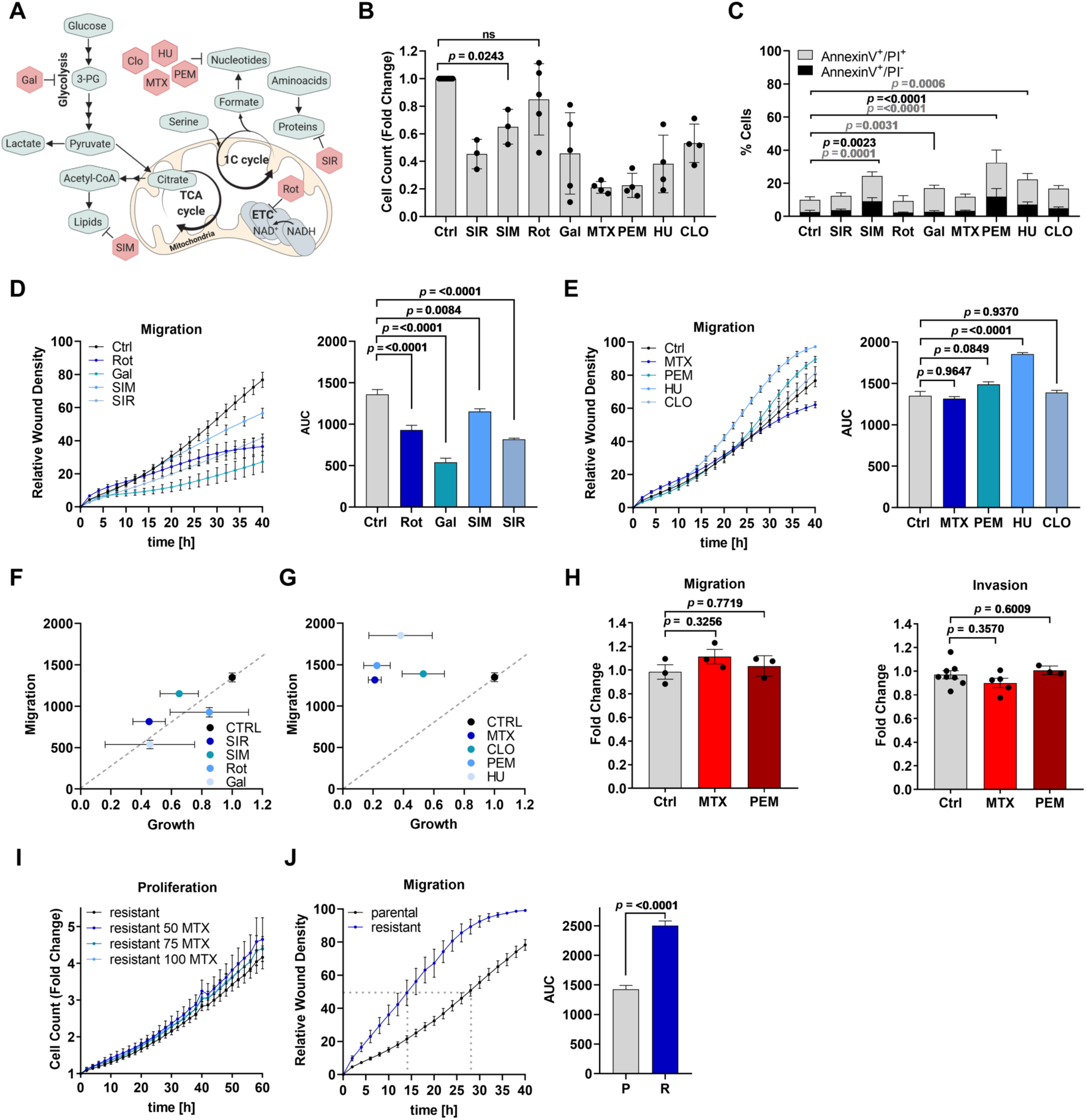
Inhibition of Major Anabolic Synthesis Routes Differentially Impacts the Motility Potential of Cancer Cells. **(A)** Synthesis routes of metabolic building blocks (lipids, nucleotides, and proteins) and pharmacologic intervention points of the selected drug panel. **(B)** MDA-MB-468 cells were counted after 48 h treatment with 100 nM Sirolimus (SIR), 1 μM Simvastatin (SIM), 50 nM Rotenone (Rot), galactose (Gal) supplementation, 50 nM Methotrexate (MTX), 1 μM Pemetrexed (PEM), 0.5 mM hydroxyurea (HU), and 100 nM Clofarabine (CLO). Each dot represents an independent experiment; mean ± SD; ordinary one-way ANOVA with Dunnett’s multiple comparisons test. Unlabeled comparisons to Ctrl are all significant with p<0.0001. **(C)** MDA-MB-468 cells were treated as in (B). Cell death was assessed by flow cytometry and AnnexinV-FITC/PI-staining; mean ± SD (n = 3 - 8). 2way ANOVA with Dunnett’s multiple comparisons test. **(D, E)** Migration of MDA-MB-468 cells upon (D) 50 nM Rot, Gal, 1 μM SIM, and 100 nM SIR or (E) 50 nM MTX, 1 μM PEM, 0.5 mM HU, and 100 nM CLO and respective area under curve (AUC); mean ± SEM (n = 3 - 12); Brown-Forsythe and Welch one-way ANOVA with Dunnett’s multiple comparisons test. **(F**,**G)** Correlation of cell migration (AUC over 40h) and proliferation (fold cell growth after 48 h). MDA-MB-468 cells were treated as in (B) and (D, E). **(H)** Migration and invasion of MDA-MB-468 cells treated for 24 h with 50 nM MTX using non-coated (migration) or ECM-Collagen-coated (invasion) Boyden chambers. Dots represent independent experiments; mean ± SEM; Brown-Forsythe and Welch one-way ANOVA with Dunnett’s multiple comparisons test. **(I)** Proliferation of MTX-resistant MDA-MB-468 cells upon the indicated concentrations of MTX [nM]; mean ± SEM (n = 4). **(J)** Migration of parental and MTX-resistant MDA-MB-468 cells and respective AUC; mean ± SEM (n = 6 - 15); unpaired t-test with Welch’s correction. Time point of 50% wound closure is indicated.

In conclusion, the perturbation of any major anabolic route results in growth repression. However, its impact on cell migration highly depends on the targeted metabolic pathway, with nucleotide synthesis being an ineffective target to abrogate cell motility. Furthermore, prolonged nucleotide synthesis inhibition in drug-resistant cells acts as a selection pressure for more motile cancer subpopulations.

### Sustained Motility upon MTX is Supported by a ROS-driven EMT Phenotype

As MTX induced the strongest growth repression without increasing cell death (**Figure 1B, C**), we used it as a tool compound to disentangle proliferation from migration to investigate the metabolic processes that support cancer cell migration. After testing multiple concentrations (**Figure S2A**), we chose a concentration of 50 nM MTX for subsequent experiments. 50 nM MTX is significantly growth-and cell cycle-arresting in MDA-MB-468 cells (**Figure 1B, S1A**) and has also been shown to be within the therapeutic window [26-29]. As this concentration of 50 nM is significantly lower compared to earlier *in vitro* studies [20, 30-33], we first set out to validate known drug effects of MTX in our system. We observed that 48 h MTX treatment of MDA-MB-468 cells resulted in altered cellular morphology with individual cells visibly increasing in cell size and exhibited an outstretched cell shape with protruding edges (**Figure 2A**) MTX-resistant MDA-MB-468 cells showed comparable morphologic alterations (**Figure S2B**). While the observed cell cycle and proliferation arrest upon MTX treatment was rapidly reversible (**Figure S2C, D**) upon MTX removal, [U-^13^C]serine tracing and metabolic flux analysis confirmed that 50 nM MTX did strongly inhibit nucleotide synthesis (**Figure 2B**). Of note, nucleotide synthesis was partially restored in MTX-resistant MDA-MB-468 cells (**Figure S2E**). Stable isotope labeling by amino acids in cell culture (SILAC) and quantitative proteomics analysis further confirmed the efficacy of the selected MTX dose as mechanistic targets of MTX such as dihydrofolate reductase (DHFR), thymidylate synthase (TYMS), and thymidine kinase (TK1) were upregulated by treatment as expected (**Figure S2F**). DHFR expression was also increased in MTX resistant cells, while FolRa expression was slightly decreased (**Figure S2G**). Moreover, superoxide dismutase (SOD) was one of the most significantly upregulated enzymes in response to treatment (**Figure S2F**), which corresponds to prior reports that depict MTX as a potent inducer of oxidative stress [30, 34, 35]. We validated these prior findings with our low dose of MTX by measuring a moderate but significant increase of mitochondrial and cytosolic reactive oxidant species (ROS) upon acute MTX treatment and in the MTX resistant cells (**Figure S2H, I, J**). In line with increased ROS levels we observed NRF2 stabilization, however, this did not result in activation of the BACH1 axis, as reported earlier for metastatic lung cancer [36, 37] (**Figure S2K, L**). Increased ROS levels upon MTX correspond to earlier reports that highlight the potential of oxidative stress as a driver of cancer cell transformation via epithelial-mesenchymal transition (EMT) [38-44]. EMT is also a known general mechanism to escape chemotherapy induced effects.

**Figure 2:**
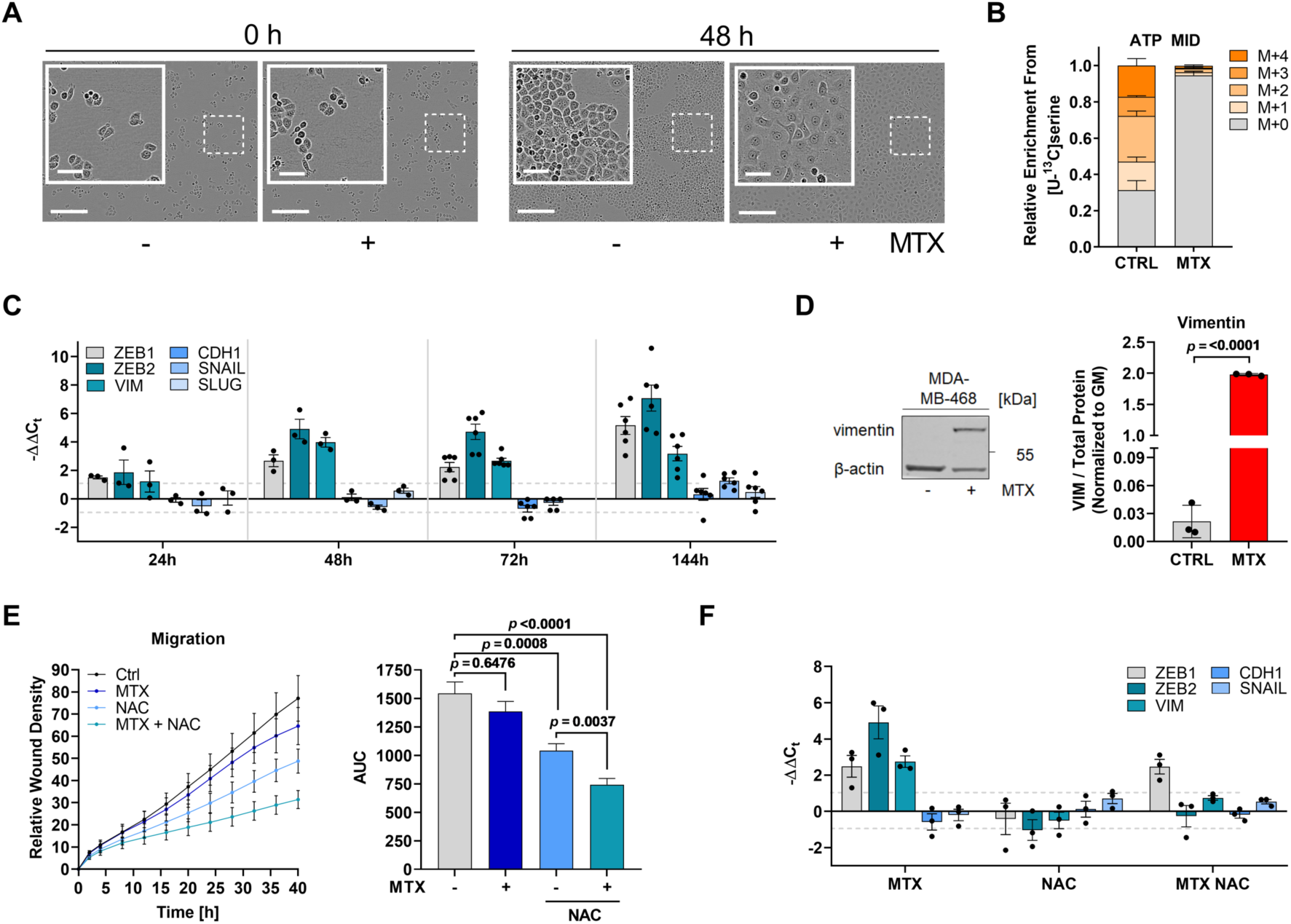
Enhanced Motility under MTX is Supported by a ROS-driven EMT Phenotype. **(A)** Morphology of MDA-MB-468 cells after 48 h 50 nM MTX. Bright-field images are representative of 3 independent experiments. Scale bars correspond to 60 and 300 μm. **(B)** Mass isotopomere distribution (MID) of intracellular ATP upon [U-^13^C]serine tracer in response to 24 h 50 nM MTX in MDA-MB-468 cells. Mean ± SEM of 5 independent experiments each measured in triplicate wells. **(C)** mRNA expression from the indicated target genes in MDA-MB-468 cells upon 50 nM MTX at the indicated time points measured by real-time RT-qPCR. Dots represent independent experiments; mean ± SEM. No data measured for *SLUG* expression at 72 h. **(D)** Protein levels of vimentin in MDA-MB-468 cells after 240 h 50 nM MTX; β-actin as loading control. Quantification of vimentin signal relative to total protein stain. Dots represent independent experiments; mean ± SD; unpaired t-test with Welch’s correction. **(E)** Migratory potential of MDA-MB-468 cells in response to 10 mM NAC and 50 nM MTX as quantification of relative wound density in IncuCyte and respective AUC. Graph shows mean ± SEM (n = 3); Brown-Forsythe and Welch ANOVA test with Games-Howell’s multiple comparisons test. **(F)** mRNA expression from the indicated target genes in MDA-MB-468 cells upon 10 mM NAC and 50 nM MTX at 72 h measured by real-time RT-qPCR. Dots represent independent experiments; mean ± SEM.

In line with previous reports, we found that ZEB1 and ZEB2 expression as well as VIM expression was upregulated in a time-dependent manner in response to acute MTX treatment and in the MTX-resistant cell line (**Figure 2C, D, S2F, M, N**). MTX-treatment also induced the expression of the collagenase MMP9 (**Figure S2O**), an enzyme which is involved in the degradation of the extracellular matrix during cancer cell invasion [45]. Treatment of cells with the antioxidant N-acetylcysteine (NAC) reduced ROS levels in MTX-treated MDA-MB-468 cells (**Figure S2P)** and resulted in a significant reduction of MTX-induced cell migration (**Figure 2E**) and EMT phenotype induction (**Figure 2F**). These findings confirm that the chosen low dose of MTX is sufficient to promote EMT in a ROS dependent manner, supporting earlier reports suggesting EMT onset in context of pulmonary fibrosis using high doses of MTX [46, 47]. Such moderate increase in cellular ROS levels after MTX might contribute to a pro-migratory stimulus that subsequently results in the observed sustained migration.

### MTX-Treated Cells Sustain High Metabolic Rates and Enhance *de novo* Serine Synthesis

In addition to the here reported MTX-specific effects on ROS and EMT, we wanted to take advantage of the MTX effect (sustained migration) to identify growth-independent metabolic liabilities that support cell migration. To that end, we profiled central carbon metabolism using [U-^13^C]glutamine and [U-^13^C]glucose tracing to monitor glycolytic activity and glutamine-or glucose-derived carbon oxidation through the TCA cycle (**Figure 3A, D**). Absolute consumption and release (CORE) rates of glutamine and glutamate were sustained in response to MTX treatment (**Figure 3B**), while the relative TCA-cycle flux of [U-^13^C]glutamine was significantly increased in response to MTX (**Figure 3C**). Absolute quantification of CORE rates of glucose and lactate also revealed sustained high glycolytic rates upon MTX treatment (**Figure 3E**). This came as a surprise, as lactate release rates were previously shown to correlate with cell growth rates [48]. In MDA-MB-468 cells, the maintained glycolytic rate and the associated lactate release at constant high levels (**Figure 3E**) indicate a constant generation of glycolysis-derived ATP even in the presence of reduced energetic demand for anabolic reactions in growth-arresting MTX conditions. Consequently, [U-^13^C]glucose distribution within the TCA-cycle was sustained by MTX treatment in MDA-MB-468 cells (**Figure 3F**) as well as in 4T1 and LN229 cells (**Figure S3A, B**). It has previously been shown that MTX inhibits oxygen consumption rates (OCR) in HCT116 cells [18]. We could replicate this finding and observed a ∼ 30 % reduction of OCR in HCT116 cells upon MTX (**Figure S3C**), while OCR in MDA-MB-468, LN229, and 4T1 cells was sustained (**Figure 3G**). This indicates cell line specific effects of MTX on OCR. Furthermore, MTX treatment significantly increased the relative flux of glucose to serine despite decreased anabolic demands for nucleotide synthesis (**Figure 3H, S3D**). Protein levels of PHGDH, PSPH, and PSAT1 that catalyze the serine *de novo* synthesis pathway from glucose were unaltered in response to MTX treatment in multiple cell lines (**Figure S3E-H**). In agreement with Diehl et al. [49] who showed that changes in NAD^+^/NADH ratio can increase PHGDH activity and in consequence serine *de novo* synthesis rates, we observed an increased NAD^+^/NADH ratio upon MTX (**Figure 3I**). As MTX is well characterized as an inhibitor of DHFR and thus 1C metabolism, it came to our surprise that MTX treatment did also increase the abundance of extracellular formate M+1 isotopologues derived from [U-^13^C]glucose through serine and the mitochondrial 1C metabolism (**Figure 3J, S3I**).

**Figure 3:**
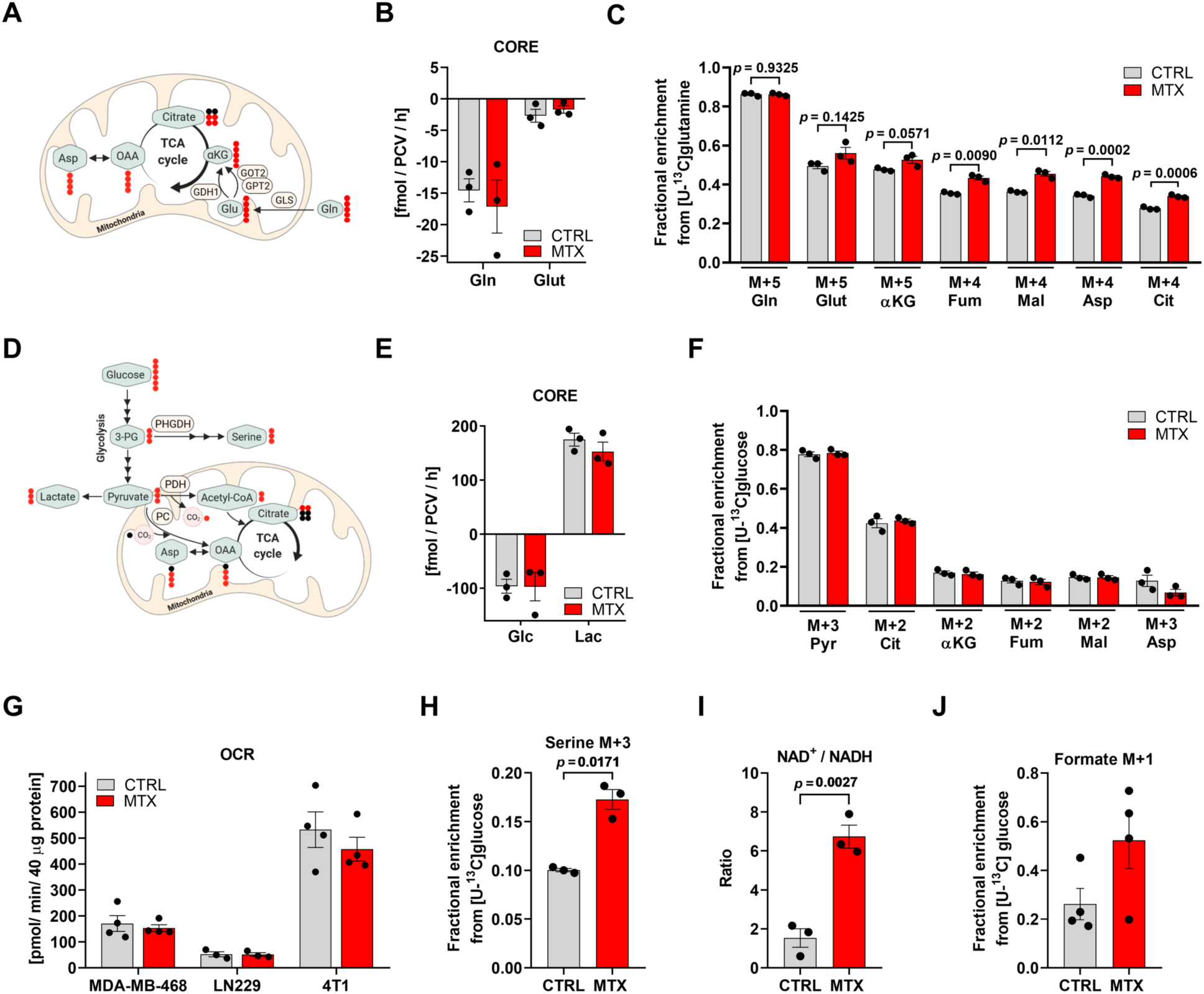
MTX-Treated Cells Sustain High Metabolic Rates and Enhance *de nov*o Serine Synthesis. **(A)** Interdependence of glycolysis and tricarboxylic acid (TCA) cycle and metabolic ^13^C label pattern from [U-^13^C]glutamine. **(B)** Absolute consumption and release (CORE) rates of glutamine (Gln) and glutamate (Glut) in the culture medium of MDA-MB-468 cells after 24 h 50 nM MTX. Dots represent individual experiments composed of triplicate wells; mean ± SEM. **(C)** Enrichment of selected isotopologues of Gln, Glut, α-ketoglutarate (αKG), fumarate (Fum), malate (Mal), aspartate (Asp), and citrate (Cit) upon [U-^13^C]glutamine tracer in response to 24 h 50 nM MTX in MDA-MB-468 cells. Dots represent independent experiments in triplicate wells; mean ± SEM; unpaired t-test with Welch’s correction. **(D)** Interdependence of glycolysis and TCA cycle and metabolic ^13^C label pattern from [U-^13^C]glucose. **(E)** Absolute CORE rates of lactate and glucose in the culture medium of MDA-MB-468 cells after 24 h 50 nM MTX. Dots represent individual experiments composed of triplicate wells; mean ± SEM. **(F)** Enrichment of selcted isotopologues of pyruvate (Pyr), Cit, αKG, Fum, Mal, and Asp upon [U-^13^C]glucose tracer in response to 24 h 50 nM MTX in MDA-MB-468 cells. Dots represent individual experiments composed of triplicate wells; mean ± SEM; unpaired t-test with Welch’s correction. **(G)** Basal cellular respiration upon 24 h 50 nM MTX in MDA-MB-468, LN229, and 4T1 cells as quantification of mitochondrial oxygen consumption rate (OCR). Dots represent individual experiments composed of six technical replicates; mean ± SEM. **(H)** Enrichment of M+3 isotopologue of serine upon [U-^13^C]glucose upon 24 h 50 nM MTX in MDA-MB-468 cells. Dots represent independent experiments in triplicate wells; mean ± SEM; unpaired t-test with Welch’s correction. **(I)** Ratio of absolute intracellular NAD^+^ to NADH in MDA-MB-468 cells upon 24 h 50 nM MTX. Dots represents individual experiments composed of triplicate wells; mean ± SEM. **(J)** M+1 isotopologue of extracellular formate in MDA-MB-468 cells from [U- ^13^C]glucose upon 24 h 50 nM MTX. Dots indicate independent experiments measured in triplicate wells; mean ± SEM.

In summary, the metabolic profiling demonstrates that, despite decreased metabolic demand for biomass production, MTX-treated cells sustain high metabolic rates that are comparable to fully proliferating cells. Additionally, and in contrast to general assumptions, cytosolic DHFR inhibition results in increased rates of serine synthesis and downstream mitochondrial formate excretion suggesting an uncoupling of mitochondrial 1C metabolism from cytosolic 1C metabolism.

### Mitochondria Protect 1C metabolism-Dependent Serine Catabolism upon Cytosolic 1C Pathway Inhibition

Our finding of increased [U-^13^C]glucose-derived M+1 formate excretion under MTX (**Figure 3J, S3I**), allows the hypothesis that mitochondrial 1C metabolism can function in an autarkic manner when cytosolic 1C metabolism is inactivated (**Figure 4A**). To corroborate this hypothesis, we employed [U-^13^C]serine tracing to monitor serine flux through 1C metabolism. We observed that cytosolic block of 1C metabolism via MTX did not reduce the exchange rates of serine, glycine and formate in MDA-MB-468 or LN229 cells (**Figure 4B, C, S4A, C**). In contrast, the formate release rate was significantly increased and the increased fraction of labeled formate proved that most of the released formate is derived from [U-^13^C]serine and not from other carbon sources that could alternatively generate formate in presence of MTX. Of note, sustained serine consumption and formate overflow via 1C metabolism upon DHFR inhibition could also be confirmed in Plasmax medium [50], which is a culture medium closer to human physiology (**Figure S4D**). Since cell dry mass composition is constituted by around 60 % of proteins [51], growth arresting conditions generally result in decreased consumption rates of proteinogenic amino acids. In fact, a general trend for decreased consumption of essential amino acids was observed (**Figure S4B**). Hence, sustained serine consumption rates upon growth arrest suggest that spared serine that is otherwise used for anabolic processes such as nucleotide synthesis, is used for alternative metabolic pathways that support cell motility and survival.

**Figure 4:**
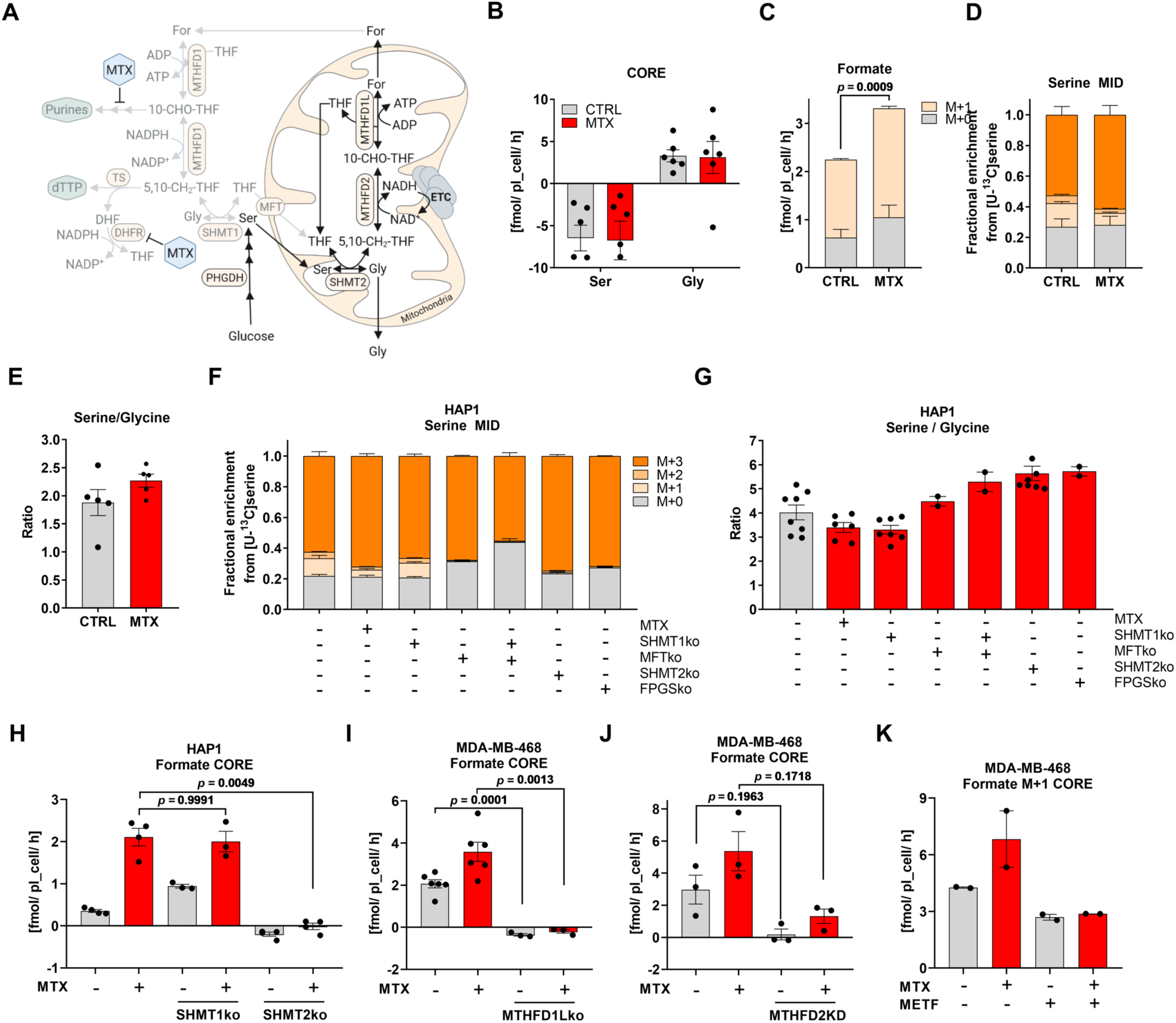
Mitochondria Protect 1C metabolism-Dependent Serine Catabolism upon Cytosolic 1C Pathway Inhibition. **(A)** Cytosolic and mitochondrial 1C metabolism and the pharmacologic intervention points of MTX. **(B)** Absolute CORE rates of serine (Ser) and glycine (Gly) from MDA-MB-468 cells upon 24 h 50 nM MTX. Dots represent the mean of an individual experiment each measured in triplicate wells; mean ± SEM. **(C)** Formate release rate upon [U-^13^C]serine in MDA-MB-468 cells after 24 h 50 nM MTX; mean ± SEM of three independent experiments each measured in triplicate wells; unpaired t-test with Welch’s correction for M+1 isotopologue. **(D)** MID of intracellular serine upon [U-^13^C]serine in MDA-MB-468 cells upon 24 h 50 nM MTX; mean ± SEM of four independent experiments each measured in triplicate wells. **(E)** Ratio of absolute intracellular serine and glycine after 24 h 50 nM MTX in MDA-MB-468 cells. Dots represent an independent experiment in triplicate wells; mean ± SEM. **(F)** MID of intracellular serine upon [U-^13^C]serine in SHMT1, MFT, SHMT2, and FPGS-depleted HAP1 cells; mean ± SEM of independent experiments each measured in triplicate wells (n = 2 – 8). **(G)** Ratio of absolute intracellular serine and glycine levels in SHMT1, MFT, SHMT2, and FPGS depleted HAP1 cells. Dots represent an independent experiment in triplicate wells; mean ± SEM. (**H**) Absolute CORE rates of formate from WT and SHMT1- or SHMT2-depleted HAP1 cells upon 24 h 50 nM MTX. Dots represent mean of an individual experiment measured in triplicate wells; mean ± SEM; Brown-Forsythe and Welch ANOVA test with Games-Howell’s multiple comparisons test. (**I**) Absolute CORE rates of formate from WT and MTHFD1L-depleted MDA-MB-468 cells upon 24 h 50 nM MTX. Dots represent mean of an individual experiment measured in triplicate wells; mean ± SEM; Brown-Forsythe and Welch ANOVA test with Games-Howell’s multiple comparisons test. (**J**) Absolute CORE rates of formate from SCR and MTHFD2 KD MDA-MB-468 cells upon 24 h 50 nM MTX. Dots represent mean of an individual experiment measured in triplicate wells; mean ± SEM; Brown-Forsythe and Welch ANOVA test with Games-Howell’s multiple comparisons test. (**K**) Absolute CORE rates of formate from MDA-MB-468 cells upon 24 h treatment with 2.5 mM Metformin and 50 nM MTX. Dots represent mean of an individual experiment measured in triplicate wells; mean ± SEM; Brown-Forsythe and Welch ANOVA test with Games-Howell’s multiple comparisons test.

Mechanistically, 1C metabolism follows a cycle in which serine can be resynthesized in the cytoplasm and mitochondrion from glycine via serine hydroxymethyltransferase 1 and 2 (SHMT1, SHMT2) (**Figure 4A**). As the SHMT reaction is highly reversible, the serine labeling pattern from [U-^13^C]serine is composed of a mix of M+1, M+2 and M+3 isotopologues, which represents the different recombination events with labeled and unlabeled glycine and formate [16, 21, 52]. Consequently, M+1 and M+2 serine isotopologues are expected to be absent upon complete 1C metabolism inhibition. Intriguingly, cytosolic DHFR inhibition with MTX neither completely abrogated serine M+1 and M+2 isotopologues upon [U-^13^C]serine nor did it significantly affect the serine to glycine level ratio in MDA-MB-468 or LN229 cells (**Figure 4D, E, S4E, F**), further supporting our hypothesis of a persistent mitochondrial 1C metabolism upon DHFR inhibition. To mechanistically disentangle cytosolic from mitochondrial 1C metabolism, we analyzed a panel of CRISPR knockouts in HAP1 cells in which either the cytosolic (SHMT1), the mitochondrial (MFT, SHMT2), or both compartments (FPGS, MFT+SHMT1) of 1C metabolism were abrogated [53]. Here, cytosolic SHMT1 KO did not eliminate intermediary M+1 and M+2 serine isotopologues from [U-^13^C]serine comparable to MTX treatment, whereas mitochondrial or combined cytosolic and mitochondrial inhibition of 1C metabolism upon MFT KO, SHMT2 KO, FPGS KO and MFT+SHMT1 KO resulted in a complete loss of M+1 and M+2 serine isotopologues (**Figure 4F**). In mutants harboring a full inhibition of 1C metabolism or mitochondrial 1C metabolism alone, we additionally observed an increase in the ratio of serine to glycine levels, whereas MTX treatment and SHMT1 KO did not substantially increase serine to glycine ratio (**Figure 4G**).

Of note the protein abundance of mitochondrial serine catabolism enzymes was not increased upon MTX. MTHFD2 abundance even decreased upon MTX, however, this did apparently not affect the rate of serine catabolism (**Figure S4G, H**).

To test if knock-out of mitochondrial serine catabolism stops formate overflow, we monitored formate release flux upon MTX in HAP1 cells deficient for SHMT1 or SHMT2. As expected, loss of SHMT2 completely abolished formate overflow, while SHMT1 KO increased formate overflow (**Figure 4H**). This experiment was then validated in MDA-MB-468 cells harboring MTHFD1L KO [20] or shRNA mediated MTHFD2 knock-down and MTHFD2 KO (**Figure S4I, K, L**). All genetic interventions resulted in a strong suppression of formate overflow during MTX treatment, further suggesting that increased formate overflow upon MTX depends on mitochondrial serine catabolism via the folate pathway (**Figure 4I, J, S4J, M**).

As an alternative approach to genetic silencing we used Metformin to target complex 1 of the electron transport chain (ETC). We have previously shown that mitochondrial formate production depends on active ETC [21, 22]. Here, we demonstrate that targeting of complex 1 with Metformin strongly suppresses mitochondrial formate production both at baseline and in presence of MTX, further supporting our finding that formate production in presence of MTX depends on mitochondrial 1C metabolism (**Figure 4K**). To proof that Metformin indeed inhibits respiration under the applied conditions, we performed Seahorse experiments to measure the expected drop in oxygen consumption rate (**Figure S4N**). We also monitored a reduction in proliferation (**Figure S4O**) and we performed [U-^13^C]glutamine tracing to monitor the expected strong increase in reductive carboxylation (**Figure S4P, Q**), a well-known phenomenon upon ETC inhibition [54, 55].

In summary, these results show that mitochondria provide a protected cellular environment that permits autarkic folate-mediated serine catabolism independent of the cytosolic folate metabolism.

### Limiting Serine availability Decreases Formate Overflow and Cancer Cell Migration

As we observed increased glucose derived serine synthesis and subsequent downstream catabolism of serine via mitochondrial 1C metabolism we investigated if interfering with serine synthesis is effective to reduce sustained cell migration upon pharmacologic inhibition of cytosolic 1C metabolism. First, to identify an effective inhibitor of serine *de novo* synthesis, we tested multiple available allosteric and one competitive inhibitor of PHGDH. Indicated by the strong abrogation of labeled serine, we observed that the competitive inhibitor BI-4916 (BI) [12] had superior efficacy compared to the allosteric inhibitors WQ-2101 (WQ) [11], NCT-502 (NCT) [9], and CBR-5884 (CBR) [15] (**Figure 5A**). Released M+1 formate from [U-^13^C]glucose was also effectively inhibited after BI treatment alone or in combination with MTX (**Figure 5B**). Additionally, routine screening for metabolic side effects revealed that treatment with the allosteric inhibitors WQ, NCT, and CBR did result in a significant reduction of mitochondrial OCR (**Figure 5C**), which was not observed with the competitive inhibitor BI (**Figure 5C**). WQ and NCT also negatively affected TCA cycle activity as characterized by a reduction of M+2 isotopologue abundance of TCA cycle associated metabolites (**Figure 5D**). In contrast to the other inhibitors, BI had no adverse effect on TCA cycle activity and proliferation rate and thus emerged as the preferred PHGDH inhibitor in all subsequent experiments (**Figure 5D, S5A-D**). Combined treatment with the specific PHGDH inhibitor BI significantly reduced MTX-mediated cell migration (**Figure 5E**). This inhibitory effect was further enhanced upon combined PHGDH inhibition and serine and glycine (S/G) starvation (**Figure 5F**). While S/G starvation alone resulted only in moderate inhibitory effects on migration of MTX-treated cells, combined treatment with PHGDH inhibition fully blunted cell migration (**Figure 5F**). Similar results were observed in 4T1 cells (**Figure S5F**). As all MTX conditions resulted in full growth arrest (**Figure S5E**), this finding indicates a role for mitochondrial 1C metabolism in cell motility that is independent of serine’s anabolic function to support biosynthetic processes.

**Figure 5:**
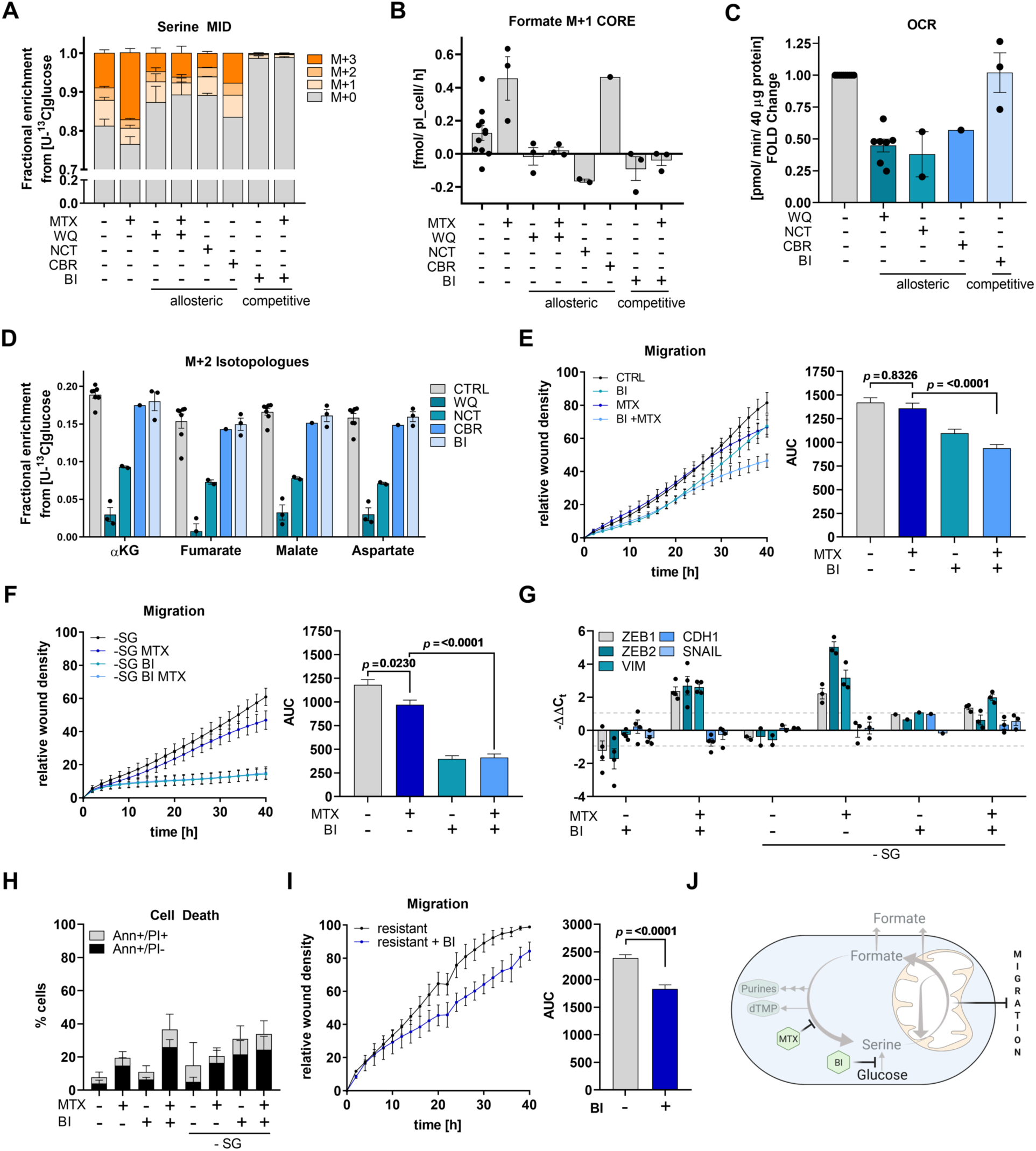
Limiting Serine availability Decreases Formate Overflow and Cancer Cell Migration. **(A)** MID of intracellular serine in MDA-MB-468 cells upon [U-^13^C]glucose and 24 h 10 μM WQ-2101, 10 μM NCT-502, 30 μM CBR-5884, 15 μM BI-4916, and 50 nM MTX; mean ± SEM of independent experiments each measured in triplicate wells (n = 1 – 9). **(B)** formate M+1 isotopologue exchange rate in MDA-MB-468 cells using [U-^13^C]glucose and treatment as in (A). Dots indicate independent experiments measured in triplicate wells; mean ± SEM. **(C)** Basal cellular respiration upon 24 h 10 μM WQ-2101, 10 μM NCT-502, 30 μM CBR-5884, and 15 μM BI-4916 in MDA-MB-468 cells as fold mitochondrial OCR compared to control. Dots represent individual experiments composed of six technical replicates; mean ± SEM. **(D)** M+2 isotopologues of selected TCA-metabolites in MDA-MB-468 cells upon [U-^13^C]glucose and treatment as in (B). Dots indicate independent experiments measured in triplicate wells; mean ± SEM. **(E**,**F)** Migration of MDA-MB-468 cells upon 50 nM MTX,15 μM BI-4916 and (F) serine- and glycine starvation and respective AUC; mean ± SEM (n = 4-5); Brown-Forsythe and Welch one-way ANOVA with Dunnett’s multiple comparisons test. **(G)** mRNA expression from indicated genes in MDA-MB-468 cells upon 72 h 50 nM MTX, 15 μM BI-4916, and serine-and glycine starvation measured by real-time RT-qPCR. Dots represent independent experiments; mean ± SEM. **(H)** Cell death induction in MDA-MB-468 cells upon 48 h 50 nM MTX, 15 μM BI-4916 and serine-and glycine starvation was assessed by flow cytometry and AnnexinV-FITC/PI-staining; mean ± SD (n = 3). 2way ANOVA with Dunnett’s multiple comparisons test. **(I)** Migration of MTX-resistant MDA-MB-468 cells upon 50 nM MTX and 15 μM BI-4916 and respective AUC; mean ± SEM (n = 4); Brown-Forsythe and Welch one-way ANOVA with Dunnett’s multiple comparisons test. **(J)** Cytosolic and mitochondrial compartments of 1C metabolism and the pharmacologic intervention points of MTX and inhibitors of PHGDH.

While neither BI treatment alone nor S/G starvation alone were sufficient to prevent the induction of the observed EMT-phenotype, combined inhibition and starvation did minimize ZEB1, ZEB2, and VIM upregulation upon DHFR inhibition (**Figure 5G**). This indicates that serine represents an underlying requirement for EMT onset upon DHFR inhibition. Importantly, S/G starvation did not further increase cell death after combined MTX and BI treatment (**Figure 5H**). Of note, we also found that BI treatment was effective to reduce the migratory capacity of MTX-resistant, pro-migratory MDA-MB-468 cells (**Figure 5I**). In summary, using MTX as a tool compound to study cell migration in absence of proliferation revealed that serine is essential to sustain full cancer cell migratory capacity (**Figure 5J**).

### Mitochondrial 1C Metabolism Supports Cell Motility and depends on mitochondrial THF availability

As withdrawal of serine can affect different aspects of cell physiology, we aimed to investigate more specifically the role of mitochondrial 1C metabolism in promoting cell migration. Analogue to the interventions presented in Figure 4I-K, we used Metformin to target the ETC and genetic interventions to target MTHFD2 or MTHFD1L both in MDA-MB-468 and 4T1 cells.

Metformin treatment of MTX exposed (growth arrested) cells reduced the migration of MDA-MB-468 cells (**Figure 6A**), suggesting that one function of the ETC is to support cell migration.

**Figure 6:**
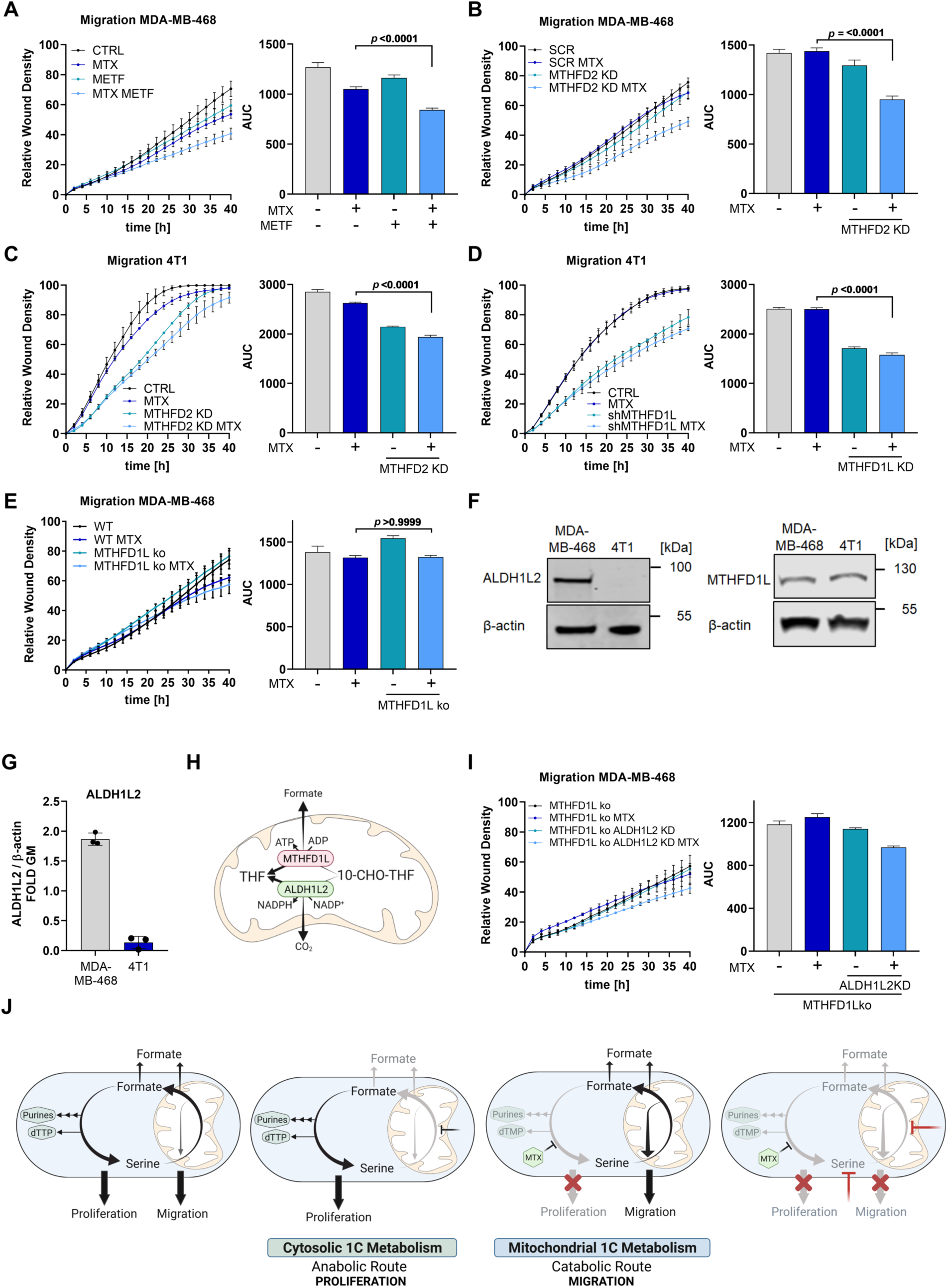
Mitochondrial 1C Metabolism Supports Cell Motility and depends on mitochondrial THF availability. **(A)** Migration of MDA-MB-468 cells upon 2.5 mM Metformin and 50 nM MTX and respective AUC; mean ± SEM (n = 3); Brown-Forsythe and Welch one-way ANOVA with Dunnett’s multiple comparisons test. **(B)** Migration of MDA-MB-468 cells upon MTHFD2 knockdown and 50 nM MTX and respective AUC; mean ± SEM (n = 4); Brown-Forsythe and Welch one-way ANOVA with Dunnett’s multiple comparisons test. (**C**) Migration of 4T1 cells upon Mthfd2 knockdown and 75 nM MTX and respective AUC; mean ± SEM (n = 3); Brown-Forsythe and Welch one-way ANOVA with Dunnett’s multiple comparisons test. (**D**) Migration of 4T1 cells upon Mthfd1l knockdown and 75 nM MTX and respective AUC; mean ± SEM (n = 3); Brown-Forsythe and Welch one-way ANOVA with Dunnett’s multiple comparisons test. (**E**) Migration of MDA-MB-468 cells upon MTHFD1L knockout and 50 nM MTX and respective AUC; mean ± SEM (n = 3); Brown-Forsythe and Welch one-way ANOVA with Dunnett’s multiple comparisons test. **(F)** ALDH1L2 and MTHFD1L protein expression in MDA-MB-468 and 4T1 cells. (**G**) Signal intensity of ALDH1L2 on Western Blot was quantified relative to β-actin as loading control. Each dot represents an independent experiment, mean ± SEM; unpaired t-test with Welch’s correction. (**H**) MTHFD1L- and ALDH1L2-dependent reactions in mitochondrial 1C metabolism. (**I**) Migration of MDA-MB-468 cells upon MTHFD1L knockout, ALDH1L2 knockdown and 50 nM MTX and respective AUC; mean ± SEM (n = 2); Brown-Forsythe and Welch one-way ANOVA with Dunnett’s multiple comparisons test. (**J**) Model summarizing the autarkic activity of mitochondrial and cytosolic 1C metabolism and their respective relevance for cell migration and cell proliferation.

Next, we investigated the role of mitochondrial serine catabolism in context of migration directly, by monitoring the migration of MTX treated *MTHFD2* or *MTHFD1L* silenced MDA-MB-468 and 4T1 cells. Of note, neither MTHFD2 nor MTHFD1L silencing reduced cell proliferation or sensitivity to the growth-arresting effects of MTX (**Figure S6A-D**). Also, cell death rates upon MTX were not affected by MTHFD2 or MTHFD1L silencing (**Figure S6E-H**). Scratch assay analysis showed that *MTHFD2* silencing markedly reduced MTX-mediated cell motility both in MDA-MB-468 and 4T1 cells (**Figure 6B, C**). As shRNA used to silence human and murine MTHFD2 were different we do not expect shRNA specific off-target effects. Nevertheless, we confirmed the shMTHFD2 KD results in MDA-MB-468 cells with MTHFD2 knock-out cells (**Figure S6I)**.

Mthfd1l silencing of MTX treated 4T1 cells also resulted in decreased migration (**Figure 6D**). Somewhat surprisingly, loss of MTHFD1L in MDA-MB-468 cells had no negative effect on the migration of MTX treated cells (**Figure 6E**). By following different hypotheses and various experiments, we realized that there is a striking difference in ALDH1L2 expression in 4T1 versus MDA-MB-468 cells, while MTHFD1L expression was comparable between these two cell lines (**Figure 6F, G**).

This observation reminded us that in MDA-MB-468 cells, ALDH1L2 can act as an alternative outlet to oxidize formyl-THF to CO^2^ thereby regenerating mitochondrial THF (in contrast to 4T1). That said, in MTHFD1L deficient cells, the ALDH1L2 status can be decisive whether THF can be regenerated or if THF will be trapped as 10-CHO-THF in the mitochondrion. Loss of free THF will inhibit the autarkic mitochondrial cycle that can otherwise act in presence of MTX (**Figure 6H**). In support of this hypothesis, we also measured increased formate release flux in 4T1 cells compared to MDA-MB468 (**Figure S6J**), suggesting that in 4T1 cells less 10-CHO-THF is oxidized via ALDH1L2 and instead catabolized through MTHFD1L.

To test this hypothesis, we targeted *ALDH1L2* in MTHFD1L KO cells and monitored migration. We found that combinatorial depletion of MTHFD1L and ALDH1L2 in MDA-MB-468 cells resulted in a reduction of migration upon MTX compared to single depletion of MTHFD1L alone. This was observed both for shRNA as well as for siRNA mediated knockdown of *ALDH1L2* (**Figure 6I, S6K**).

In conclusion, these results reveal a growth-independent role of mitochondrial 1C metabolism for cancer cell motility. Interestingly, the pool of free THF might be relevant for the functionality of the autarkic mitochondrial 1C cycle and ALDH1L2 can serve as an alternative outlet to regenerate THF when MTHFD1L is missing (summarized in **Figure 6J**).

### Genetic Targeting of Mitochondrial 1C Metabolism Reduces Metastasis Formation *in vivo*

According to our model, we predict that sole inhibition of mitochondrial 1C metabolism might be sufficient and effective to reduce cancer cell escape from the primary tumor and subsequent metastasis formation. To test this hypothesis, we employed the 4T1 cells with *Mthfd1l* KD to study the metastatic cascade in orthotopic tumors in vivo.

To evaluate first if Mthfd1l KD cells are more prone to oxidative stress, which could impact metastatic dissemination *in vivo* [56], we measured ROS levels in MTHFD1L silenced MDA-MB-468 and 4T1 cells at baseline and after MTX treatment. At baseline, ROS levels between MTHDF1L silenced and control cells were unchanged. MTX treatment induced ROS levels to comparable extends and 500 μM of H^2^O^2^ treatment did induce cell death rate comparable to Ctrl cells (**Figure S7A-C**). In summary, these experiments do not indicate that loss of MTHFD1L results in increased cellular ROS levels or increased susceptibility to oxidative insults. We then injected 4T1 cells with or without silenced Mthfd1l into the mammary fat pad of immunocompetent BALB/c mice to form orthotopic tumors (**Figure 7A**). Primary tumor growth was monitored over 6 weeks. In concordance with the *in vitro* data, primary tumor growth was not affected and tumor weight at endpoint was not different compared to Ctrl conditions (**Figure 7B, C**). Knock-down of Mthfd1l in the primary tumors was confirmed by Western Blot (**Figure 7D, S7D-F**). In line with our hypothesis, the number of macroscopic lung metastases was significantly reduced in mice that were carrying tumors deficient in Mthfd1l (**Figure 7E**). While all mice that were injected with Mthfd1l-expressing 4T1 cells exhibited lung metastases, over 50% of mice injected with *Mthfd1l* KD cells did not show macroscopic metastatic lesions. Of note, expression of EMT related genes in the primary tumor was unchanged between the two groups (**Figure S7G**). H&E staining of lungs and quantification of the metastatic area per lung confirmed reduced metastasis in Mthfd1l deficient tumors (**Figure 7F, G)**.

**Figure 7:**
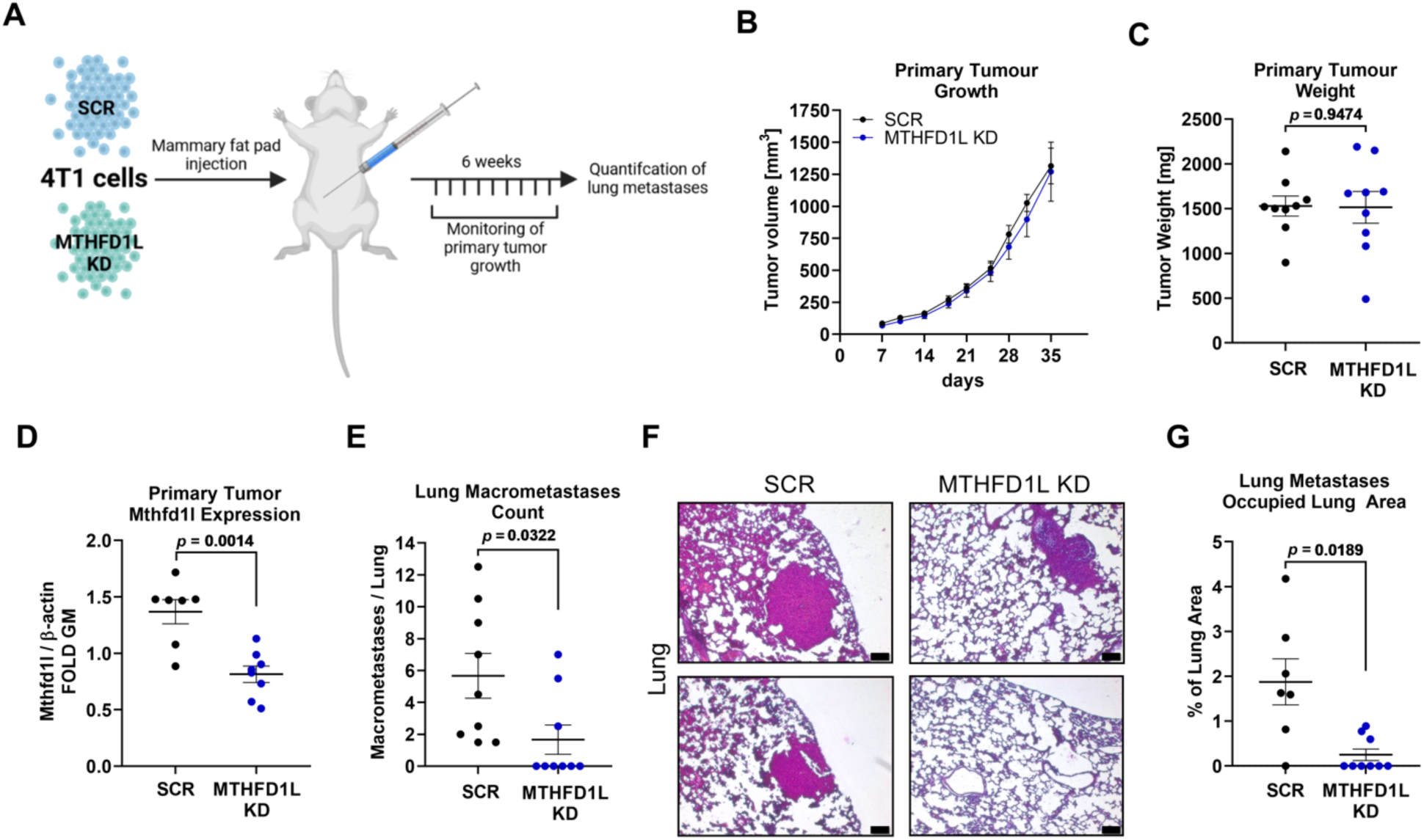
Genetic Targeting of Mitochondrial 1C Metabolism Reduces Metastasis Formation *in vivo*. **(A)** 4T1 breast cancer cells transfected with non-targeting control (SCR, n=9) and 4T1 cells with KD of *Mthfd1l* (n=9) were injected into the mammary fat pads of immunocompetent female Balb/c mice. Primary tumor growth was monitored and lung metastasis formation was evaluated at end-point. **(B)** Primary tumor size was measured and mean tumor volume for each group was calculated; mean ± SEM. **(C)** Primary tumor weight was measured at the end-point. Dots indicate individual animals; mean ± SEM; unpaired t-test with Welch’s correction. **(D)** Primary tumor tissue was analyzed for Mthfd1l protein expression by Western Blot and quantified relative to β-actin as loading control. Dots indicate individual animals; mean ± SEM; unpaired t-test with Welch’s correction. Protein lysates from tumors with significantly low Mthfd1l and β-actin expression relative to total protein were judged to be not pure tumor lysate and discarded from analysis (see Figure S7D-F for details). (**E**) Macroscopic lung metastases were counted and are depicted as number of metastases per lung. Dots indicate individual animals; mean ± SEM; unpaired t-test with Welch’s correction. (**F**) Lung tissue was embedded in paraffin and stained with H&E. 2 representative images per group are presented to show microscopic lung metastasis; scale bar corresponds to 100 μm. **(G)** metastatic area per lung was quantified and is expressed as % area of lung tissue. Dots indicate individual animals; mean ± SEM; unpaired t-test with Welch’s correction.

These *in vivo* findings further strengthen the key relevance of mitochondrial 1C metabolism as a cornerstone to support cancer cell migration independent of primary tumor growth and cancer cell proliferation rates.

## Discussion

In this study, we uncover a growth-independent function of mitochondrial serine catabolism to drive cancer cell motility. Importantly, by using MTX as a tool compound to study migration independent of proliferation, we observed that upon inhibition of cytosolic 1C metabolism, mitochondria sustain an autarkic 1C metabolism that is sufficient to sustain full migratory capacity.

Such compartmentalization of 1C metabolism during MTX treatment is feasible because release of the 1C unit from THF in the mitochondrial MTHFD1L reaction is, in contrast to cytosolic thymidylate synthetase reaction, not oxidizing THF to DHF. Therefore, the mitochondrial THF pool is sustained to allow SHMT2-dependent mitochondrial serine catabolism, while cytosolic 1C metabolism enzymes are inhibited due to accumulating DHF species. Our finding that ALDH1L2 can serve as an alternative source for THF regeneration in MTHFD1L deficient cells, supports the notion that mitochondria can maintain an independent THF pool that is required for an autarkic mitochondrial 1C metabolism as long as there is THF available that can accept the hydroxymethyl group coming in the first (SHMT2 dependent) step of serine catabolism.

An autarkic mitochondrial 1C metabolism is further supported by a chemical modification of folate species that alters their transport activities across the mitochondrial membrane. For maximal biologic activity, folate species need to be polyglutamated by folylpolyglutamate synthase (FPGS) and the resulting polyglutamate species were shown to be only poorly transported across the mitochondrial membrane [57, 58]. Consequently, polyglutamated folate species can be chemically trapped within the mitochondria to sustain the mitochondrial 1C cycle. Especially, upon growth arrest when cells can retain their mitochondrial content and don’t need to amplify their content as it is required during proliferation. Additionally, there is evidence that transport activity of the mitochondrial folate transporter is limited to reduced folates [59]. This indicates a limited transport of MTX and folic acid itself across the mitochondrial membrane. Taken together, such compartmentalization of 1C metabolism and autarkic function of mitochondrial 1C metabolism emerges as a selective advantage in cells upon perturbation of cytosolic 1C metabolism and indicates a function of the mitochondrial 1C metabolism pathway to support cell motility.

While our data demonstrate that mitochondrial serine catabolism is needed to allow full migratory capacity, the specific biochemical output conferring this function still needs to be determined in more detail. Mitochondrial serine catabolism can generate NADH, NADPH, ATP, glycine, CO^2^, methylene-and formyl-THF and formate. The plasticity of this pathway allows tailoring the catabolism towards current cellular needs [38]. Therefore, it can be possible that serine is differentially catabolized along the metastatic cascade in a context dependent manner. We have previously shown that exogenous formate promotes cancer cell invasion [22] and various cellular stress situations such as energy stress [21] or MTX treatment (this study) can increase the rate of formate overflow which results in increased extracellular formate concentrations. It is therefore tempting to speculate that increasing formate concentrations in the tumor microenvironment promote a more invasive phenotype of stressed cancer cells. In more general terms, it could be possible that mitochondrial serine catabolism can selectively promote cell motility in response to extrinsic or intrinsic stress stimuli such as growth inhibition or nutrient deprivation. On the other hand, MDA-MB-468 MTHFD1L KO cells are also formate overflow negative but do not show reduced migration. As ROS levels are neither increased in MDA-MB-468 nor in 4T1 MTHFD1L deficient cells, we do not think that the observed migration effects in this study are ROS dependent. Given previous work [22] and the current study, there might also be differences when comparing migration with invasion. Ongoing work in our lab will help clarifying these questions in the future. To do so, it will be of particular importance to further disentangle effects arising from exogenous formate from effects depending on endogenous serine catabolism. Independent of these yet unknown details, we remain with the conclusion that mitochondrial serine catabolism contributes to cancer cell migration and metastasis independent of its known anabolic functions.

Finally, we would like to note that although we mainly use MTX as a tool compound to separate proliferative from migratory processes and to inhibit cytosolic 1C metabolism, our results have also some implications in regard to MTX as an anchor drug in chronic treatment of autoimmune diseases. Intracellular erythrocyte and mean plasma concentration of MTX were reported to be in a comparable, even slightly higher, nM range to our chosen drug concentration [60, 61]. Hence, our findings on mitochondrial 1C metabolism might justify future investigations in cancer patients and patients that undergo chronic MTX therapy for an arthritic comorbidity.

## Supporting information

Supplemental Figures Kiweler et al_012022

## Acknowledgment

We thank Fabien Rodriguez (Letellier lab) for technical support during the tissue analysis of the *in vivo* experiment. We are grateful to Lewis Cantley (Weill Cornell Medical College) for providing MDA-MB-468 cells with MTHFD1L KO [20] and to Patel’s laboratory (Cambridge, UK) for providing various HAP1 KO cells [53] We thank Clément Thomas (LIH, Luxembourg) for providing 4T1 cells and Saverio Tardito (Cancer Research UK Beatson Institute, Glasgow, UK) for providing Plasmax medium.

We would like to thank: the LCSB Metabolomics Platform, especially Xiangyi Dong and Floriane Vanhalle, for providing technical and analytical support; the National Cytometry Platform (Quantitative Biology Unit, LIH) and especially Thomas Cerutti for support with flow cytometric analyses; Magretta Adiamah for her diligent proofreading of the manuscript; all our collaboration partners for fruitful discussions and constructive feedback. All graphical figures were produced with BioRender.com.

J.M. is supported by the FNR-ATTRACT program (A18/BM/11809970), J.M. and L.H. are supported by the FNR-PRIDE NEXTIMMUNE (PRIDE/11012546) and J.M. and K.E. by the FNR-PRIDE i2Tron (PRIDE19/14254520) program. N.K. is supported by the LIH Career Launchpad program (Legs Baertz) and by a DFG fellowship (KI 2508/1-1). D.B. is supported by the FNR-ATTRACT program (A14/BM/7632103), the FNR-CORE (C18/BM/12691266), i2Tron (PRIDE19/14254520) and the FNR-RIKEN (TregBar/11228353) grant. D.B. and L.S.B. are funded by the FNR-PRIDE (PRIDE/11012546/NEXTIMMUNE) scheme. E.L. is supported by the FNR-CORE program (C16/BM/11282028 and C20/BM/14591557), by a Proof of Concept FNR grant (PoC/18/12554295), a PRIDE17/11823097 and by i2Tron (PRIDE19/14254520). G.D. and F.X. are supported by FNR-CORE (C17/BM/11642138). B.J. and M.Z.N. are supported by Luxembourg National Research Fund (C18/BM/12670304/COMBATIC) and FNRS Televie grant (7.4579.20).

## Author Contributions

Conceptualization, J.M., N.K.; Methodology, J.M., N.K., C.J., M.Z.N., B.J., E.L., D.B., G.D., V.I.P.; Software, A.V., F.X.; Validation, N.K., J.M., L.N., C.D.; Formal Analysis, N.K., C.D., L.N., M.B., L.S.B., F.X., V.I.P., M.S., C.J.; Investigation, N.K., C.D., L.N., M.B., L.S.B., F.X., V.I.P., M.S., E.L.; Resources, C.J., A.V., J.M., E.L.; Data Curation, N.K., J.M., F.X., A.V.; Writing – Original Draft, N.K., J.M.; Writing – Review & Editing, All Authors; Visualization, N.K., J.M., L.S.B., F.X., V.I.P.; Supervision, J.M., D.B., A.V., E.L., G.D.; Project Administration, J.M., Funding Acquisition, J.M., D.B., E.L., A.V., G.D..

## Materials & Methods

### Chemicals

Methotrexate, rotenone, PKUMDL-WQ-2101, galactose, hydroxyurea, NAC, metformin and fluorouracil were purchased from Sigma Aldrich. Clofarabine, pemetrexed disodium hydrate, sirolimus were purchased from Bio Connect. Simvastatin was purchased from Sanbio. CBR-5884 and BI-4916 were purchased from MedChemExpress. NCT-502 was purchased from ApeXBio.

### Cell Culture

All cell lines were cultured in Dulbecco’s modified Eagle’s medium (DMEM) without phenol red, glucose, and glutamine (Thermo Fisher Scientific) and supplemented with 2 mM glutamine, 17 mM glucose, and 10 % fetal bovine serum (FBS) at 37°C and 5 % CO^2^. For galactose treatment, supplemented glucose was replaced by 17 mM galactose at the beginning of the experiment. For serine/glycine starvation experiments cells were kept in MEM (Thermo Fisher Scientific) supplemented with or without 400 μM glycine and serine, 2 mM glutamine, 17 mM glucose and 10 % dialyzedFBS.

MDA-MB-468 and LN229 were obtained from ATCC. HCT-116 was obtained from the group of E. Letellier (LSRU, University of Luxembourg), and 4T1 from C. Thomas (LIH, Luxembourg). HAP1 cells were obtained from Patel’s laboratory [53] and cultured in IMDM medium supplemented with 10 % FBS. MTX-resistant MDA-MB-468 cells were generated through prolonged culturing of MDA-MB-468 cells with 50 nM MTX for 2 months until cells started to regrowth under MTX. All cell lines were routinely tested for mycoplasma contamination. MDA-MB-468 MTHFD1L CRISPR/Cas9 KO cells were obtained from [20].

### Metabolic Characterization

For metabolic characterization of cell lines, previously established protocols for absolute quantification of exchange fluxes and intracellular fluxes of one-carbon metabolism were applied [21].

### Stable Isotope Tracing and Metabolite Extraction

Stable isotope tracing experiments with [U-^13^C]-glucose tracer or [U-^13^C]-glutamine tracer (Cambridge Isotope Laboratories, CLM-1396) were performed in DMEM supplemented with 2 mM glutamine, 17 mM glucose tracer, and 10 % FBS. Stable isotope tracing experiments with [U-^13^C]-serine tracer (Cambridge Isotope Laboratories) were performed in MEM (Thermo Fisher Scientific) supplemented with 400 μM glycine, 2 mM glutamine, 17 mM glucose, 400 μM serine tracer, and 10 % FBS. To allow for adaptation, cells were cultivated in MEM for at least one passage prior to [U-^13^C]-serine tracer experiments. 150,000 to 200,000 cells were seeded in 12-well plates in triplicates for each experimental condition. Identical triplicate wells were seeded to allow for cell count and cell volume determination (to calculate the packed cell volume (PCV)) at the start and end of each tracing experiment. The day after seeding, growth medium was replaced by tracer medium and cells were cultured for 24 h. In parallel, 3 wells per condition were counted to assess starting PCV. After 24 h, triplicate wells were counted to assess PCV at the end of the experiment and one set of triplicates was used for subsequent metabolite extraction. Medium of these triplicates were collected and analyzed for exchange rates. To determine the basal medium composition for the subsequent calculation of exchange rates, identical medium was incubated in triplicates in empty 12 wells throughout the experiment and analyzed in parallel to the medium samples. Collected medium samples were centrifuged at 300 g for 5 min. Supernatant was collected and stored at -20°C until extraction of metabolites. Formate extraction, derivatization, and quantification as well as metabolite extraction for LC-MS analysis was performed as described in [21]. For metabolite extraction for GC-MS analysis after [U-^13^C]-glucose tracing and [U-^13^C]-glutamine tracing, cells were washed with cold 0.9 % NaCl solution. 400 μl ice-cold MeOH/H^2^O^MQ^ [(ratio, 1:1) containing the internal standards pentanedioc-d6 acid and [U-^13^C]-ribitol at a final concentration of 1 μg/ml and Tridecanoid-d25 acid at a final concentration of 5 μg/ml] was added to each well. Plates were incubated for 5 min at 4°C on a rocking shaker. Supernatant was collected, mixed with 200 μl CHCl^3^, and centrifuged for 5 min at 13,000 g at 4°C. Upper polar phase was collected and stored at -20°C for subsequent MS analysis of polar metabolites.

### GC-MS Measurements

#### Analysis of Formate Release Rates

Formate derivatization in the culture medium was performed using MCF derivatization as described in [21]. GC-MS analysis was performed using an Agilent 7890A GC coupled to an Agilent 5975C inert XL Mass Selective Detector (Agilent Technologies). A sample volume of 1 μl was injected into a Split/Splitless inlet, operating in split mode (20:1) at 270 °C. The gas chromatograph was equipped with a 30 m (I.D. 250 μm, film 0.25 μm) DB-5MS capillary column (Agilent J&W GC Column, 122-5532G). Helium was used as carrier gas with a constant flow rate of 1.4 ml/min. GC oven temperature was held at 80 °C for 1 min and increased to 130 °C at 10 °C/min followed by a post run time of 4 min at 280 °C. Total run time was 15 min. Transfer line temperature was set to 280 °C. Mass selective detector (MSD) was operating under electron ionization at 70 eV. MS source was held at 230 °C and the quadrupole at 150 °C. For precise quantification, measurements were performed in selected ion monitoring mode. Target ions (*m/z*) and dwell times are shown in Table S2. GC-MS chromatograms were processed using Agilent MassHunter Quantitative Analysis for GC-MS, Version B.08.00. Final determination of release rates was performed as described in [21].

#### Determination of mass isotopomere distribution (MID) of Intracellular TCA Cycle Metabolites following Stable Isotope Tracing

Polar metabolites were derivatized for 90 min at 45 °C with 20 μl of methoxyamine (c = 20 mg/ml) in pyridine under continuous shaking and subsequently for 90 min at 45 °C with 20 μl of MTBSTFA w/ 1% TBDMCS. GC-MS analysis was performed using an Agilent 7890B GC coupled to an Agilent 5977A Mass Selective Detector (Agilent Technologies). A sample volume of 1 μl was injected into a Split/Splitless inlet, operating in splitless mode at 270 °C. Gas chromatograph was equipped with a 30 m (I.D. 250 μm, film 0.25 μm) ZB-35MS capillary column with 5 m guard column (Phenomenex). Helium was used as carrier gas with a constant flow rate of 1.2 ml/min. GC oven temperature was held at 100 °C for 2 min and increased to 300 °C at 10 °C/min and held for 4 min. Total run time was 26 min. Transfer line temperature was set to 280 °C. Mass selective detector (MSD) was operating under electron ionization at 70 eV. MS source was held at 230 °C and the quadrupole at 150 °C. For precise quantification of the MID, measurements were performed in selected ion monitoring mode. Target ions (*m/z*) and dwell times are shown in Table S3.

The MetaboliteDetector software package (Version 3.220180913) was used for mass spectrometric data post processing, quantification, MID calculations, correction of natural isotope abundance, and determinations of fractional carbon contributions [62].

#### Analysis of Medium Exchange Rates

Polar metabolites of the culture medium were derivatized for 90 min at 45 °C with 20 μl of methoxyamine (c = 20 mg/ml) in pyridine under continuous shaking and subsequently for 90 min at 45 °C with 20 μl of MSTFA. GC-MS analysis was performed using an Agilent 7890B GC coupled to an Agilent 5977A Mass Selective Detector (Agilent Technologies). A sample volume of 1 μl was injected into a Split/Splitless inlet, operating in splitless mode at 270 °C. Gas chromatograph was equipped with a 30 m (I.D. 250 μm, film 0.25 μm) ZB-35MS capillary column with 5 m guard column (Phenomenex). Helium was used as carrier gas with a constant flow rate of 1.2 ml/min. GC oven temperature program: 90° C for 1 min, 9° C/min to 270° C, 25° C/min to 320° C and held for 7 min. Total run time was 30 min. Transfer line temperature was set to 280° C. MSD was operated under electron ionization at 70 eV. MS source was held at 230° C and the quadrupole at 150° C. Full scan mass spectra were acquired between m/z 70 and 700.

The MetaboliteDetector software package (Version 3.220180913) was used for quantification. Briefly, peak areas of all isotopologues of defined quantification ions were summed up and divided by the peak area of the internal standard for normalization. In addition, a calibration curve was prepared to calculate absolute concentrations. Absolute uptake and release rates were calculated as described in [21]

### LC-MS Measurements

Untargeted LC-MS analysis was carried out as previously described in [21].

#### Nucleotide and NAD/NADH Analysis

The following analytical conditions are based on a protocol from [21]. Metabolite analyses were performed using a Thermo Vanquish Flex Quaternary LC coupled to a Thermo Q Exactive HF mass spectrometer. Chromatography was carried out with a SeQuant ZIC-pHILIC 5μm polymer (150 × 2.1 mm) column connected to the corresponding SeQuant ZIC-pHILIC Guard (20 × 2.1 mm) pre-column. Column temperature was maintained at 45 °C. The flow rate was set to 0.2 mL/min and the mobile phases consisted of 20 mmol/L ammonium carbonate in water, pH 9.2 (Eluent A) and Acetonitrile (Eluent B). The gradient was: 0 min, 80% B; 2 min, 80% B; 17 min, 20% B; 18 min 20% B; 19 min 80 % B; 20 min 80% B (0.4 mL/min); 24 min 80% B (0.4 mL/min); 24.5 min 80% B. The injection volume was 5 μL. All MS experiments were performed using electrospray ionization with polarity switching enabled (+ESI/-ESI). The source parameters were applied as follows: sheath gas flow rate, 25; aux gas flow rate, 15; sweep gas flow rate, 0; spray voltage, 4.5 kV (+) / 3.5 kV (–); capillary temperature, 325 °C; S-lense RF level, 50; aux gas heater temperature, 50 °C. The Orbitrap mass analyzer was operated at a resolving power of 30,000 in full-scan mode (scan range: *m/z* 75‶1000; automatic gain control target: 1e6; maximum injection time: 250 ms). Data were acquired with Thermo Xcalibur software (Version 4.3.73.11) and analyzed with TraceFinder (Version 4.1). Subsequent data analysis for normalization and natural isotope subtraction were performed using in house scripts as in [21].

### Western Blot

Total cell lysates from cells cultured in vitro were prepared by 30 min incubation of cell pellets on ice in cell lysis buffer (150 mM NaCl, 1 mM EDTA, 50 mM Tris-HCl, 1% NP-40). Protein lysates from tumor tissue were obtained by lysing 10 mg tumor tissue pieces in 150 μl cell lysis buffer. Homogenization of tumor tissue lysates was achieved with TissueLyser II (Quiagen) using 5 mm metal beads. Lysis efficiency was maximized by sonification. Lysis solution was centrifuged at 13,000 g for 15 min at 4°C and supernatant was collected and stored at -80°C. Protein concentration was determined by Bradford assay. 15 – 30 μg of total protein were loaded on NuPAGE 4-12% Bis-Tris gels (Life Technologies) or RunBlue 4-12% Bis-Tris gels (Westburg) using 4x NuPage LDS Sample buffer (Thermo Fisher Scientific) supplemented with 10 mM DTT (Sigma Aldrich) and blotted on nitrocellulose membrane according to standard protocols. Membranes were stained with REVERT staining solution (LI-COR) and analyzed for total protein abundance. Subsequently, membranes were blocked with Odyssey TBS blocking buffer (LI-COR) or 5% milk-powder in TBST and incubated with the indicated primary antibodies over night at 4°C. Incubation with secondary antibody occurred for 2 h at RT. Detection was performed with the Odyssey CLx Infrared Imaging System (LI-COR). ImageStudioLite Software Vers.5.2 (LI-COR) was used for image analysis. Antibodies used for Western blot analysis in this study: MTHFD1L (16113-1-AP) from Proteintech; vimentin (3390), β-actin (3700), NRF2 (12721) and MTHFD2 (41377) from Cell Signaling Technology; PHGDH (HPA021241), PSAT1 (HPA042924), PSPH (HPA020376), SHMT1 (HPA023314), and SHMT2 (HPA020549) from Sigma Aldrich; LaminB (ab16048) from Abcam; IRDye 680RD Goat Anti-Mouse IgG (H+L) and IRDye 800CW Donkey Anti-Rabbit IgG (H+L) from LI-COR.

### Poly-L-Lysine Coating

Poly-L-lysine (P1274) was purchased from Sigma-Aldrich and reconstituted at 500 μg/ml in H^2^O^MQ^. Wells were coated with PLL prior to Seahorse measurement and Scratch assay. To this end, PLL solution was diluted 1:20 in H^2^O^MQ^ and added to the plates at least 1 h prior to seeding. Following incubation with PLL at 37°C and prior to cell seeding, plates were washed twice with H^2^O^MQ^ and allowed to air-dry.

### Seahorse Measurements

The day prior to measurement, 40,000 cells were seeded on poly-L-lysine coated plates and treated the subsequent day as indicated. XF96 Extracellular Flux Analyzer (Seahorse Bioscience) was used to measure basal OCR following manufacturer’s instructions. OCR was normalized to the protein concentration or cell number in the wells following the protocol described in [63] using Bradford assay.

### Flow Cytometric Analysis of Cell Cycle Distribution and Cell Death

200,000 cells were seeded in 2 ml DMEM and treated the subsequent day as indicated. After incubation, medium was collected and cells were washed with PBS. PBS fraction was collected and cells were detached with trypsin. Detached cells were collected in DMEM. The combined, collected solutions were centrifuged and pellet was washed with PBS.

#### Cell Cycle Distribution

Centrifugation yielded a pellet that was resuspended in 100 μl PBS and fixed with ice-cold 80% EtOH. Fixed cells were stored at -20°C for at least 1 h and maximum 5 days prior to measurement. Cells were centrifuged, pellet was incubated for 1 h in 200 μl RNAse A in PBS (30 μg/ml) at RT. Immediately prior to measurement, 98 μl propidium iodide (PI) in PBS (50 μg/ml) was added. Flow cytometric analysis was performed using BD FACSCanto and BD FACSDiva software. Analysis was performed in FlowJo.

#### Cell Death Analysis

Pellet following centrifugation was resuspended in 50 μl AnnexinV-FITC staining solution (5% AnnexinV-FITC in AnnexinV binding buffer (10 mM HEPES pH 7.4, 140 mM NaCl, 2.5 mM CaCl^2^, 0.1% BSA in ^dd^H2O)) and incubated for 15 min on ice in the dark. 450 μl PI-staining solution (1.1 μg/ml PI in AnnexinV binding buffer) was added immediately prior to measurement using BD FACSCanto and BD FACSDiva software (for wild-type cells) or NovoCyte Quanteon (for GFP-positive transfected cells). Analysis was performed in FlowJo.

### Flow Cytometric Analysis of ROS Levels

250,000 MDA-MB-468 cells were seeded in 2 ml medium and treated the subsequent day as indicated. Following incubation, cells were detached with trypsin, centrifuged at 350 g for 5 min and washed with warm DMEM. Cells were stained in 100 μl DMEM supplemented with 1:2,000 DAPI and 1:500 DCFDA for 30 min at 37°C. Following incubation, samples were centrifuged at 350 g for 5 min and washed with PBS. Following centrifugation at 350 g for 5 min, cells were resuspended in 100 μl PBS and measured using the BD LSRFortessa− system and BD FACSDiva software. Data analysis was performed using FlowJo software.

### Cell Proliferation

Cell proliferation was determined using the IncuCyte® Live-Cell Analysis system (Essen Bioscience). Cell proliferation was either determined as measurement of cell density (confluence) or by quantification of red nuclei using the IncuCyte ® NucLight Rapid Red Reagent (Essen Bioscience). Staining was performed according to manufacturer’s instructions. Alternatively, viable cell number was determined by trypan blue staining and automatic counting using a Countess™ Cell Counting Chamber Slide (Thermo Fisher).

### Migration Assay (Wound Closure)

Cell migration was determined using the IncuCyte® Scratch Wound Assay system for 96 well plates (Essen Bioscience). 96 well image lock plates (Essen Bioscience) were PLL coated prior to seeding of 40,000 cells per well. Wound application was performed 24 h post seeding using the 96-pin IncuCyte WoundMaker tool (Essen Bioscience) and indicated treatments were applied simultaneously. Measurement was performed in technical replicates (n = 4 – 8) and repeated as independent, biological experiments as stated for the respective experiment in figure legends.

### Invasion and Migration Assay (Transwell Assay)

Invasion and migration was analyzed by Boyden chamber assay as described in Meiser et al., 2018 [22]. Transwells were coated with ECM-Collagen for invasion assays and left un-coated for migration assay. Treatment and migration/invasion was carried out for 18 h to 24 h.

### RNA Extraction, cDNA Synthesis, and qPCR Analysis

Total RNA was extracted from cell culture dishes following manufacturer instructions of RNeasy Mini Kit (Qiagen, 74104) or NucleoSpin RNA Kit (Machery&Nagel, 740955). For RNA extracts from tumor tissue, 10 mg tissue were homogenized in 350 μL RNA using 5 mm metal beads in TissueLyser II (Quiagen). RNA was transcribed to cDNA with High-capacity cDNA Reverse Transcription Kit (ThermoFisher, 4368814). qPCR was performed from 20 ng cDNA per sample using Fast SYBR™ Green Master Mix (ThermoFisher, 4385612). All samples were analyzed in technical duplicates or triplicates. qPCR using Fast SYBR Green was conducted at 95°C for 20 s, 40 cycles of 95°C for 1 s and 60°C for 20 s using the QuantStudio 5 Real-Time PCR System (Applied Biosciences, ThermoFisher Scientific). Specificity was verified by melt curve analysis. Relative quantification of each mRNA was done by QuantStudio Design&Analysis v1.5.1 software (Applied Biosciences, ThermoFisher Scientific) and comparative Ct method. All expression data was normalized to two housekeeping genes (*GAPDH* and *CycloA*). All human primers used in this study are listed in the table below:

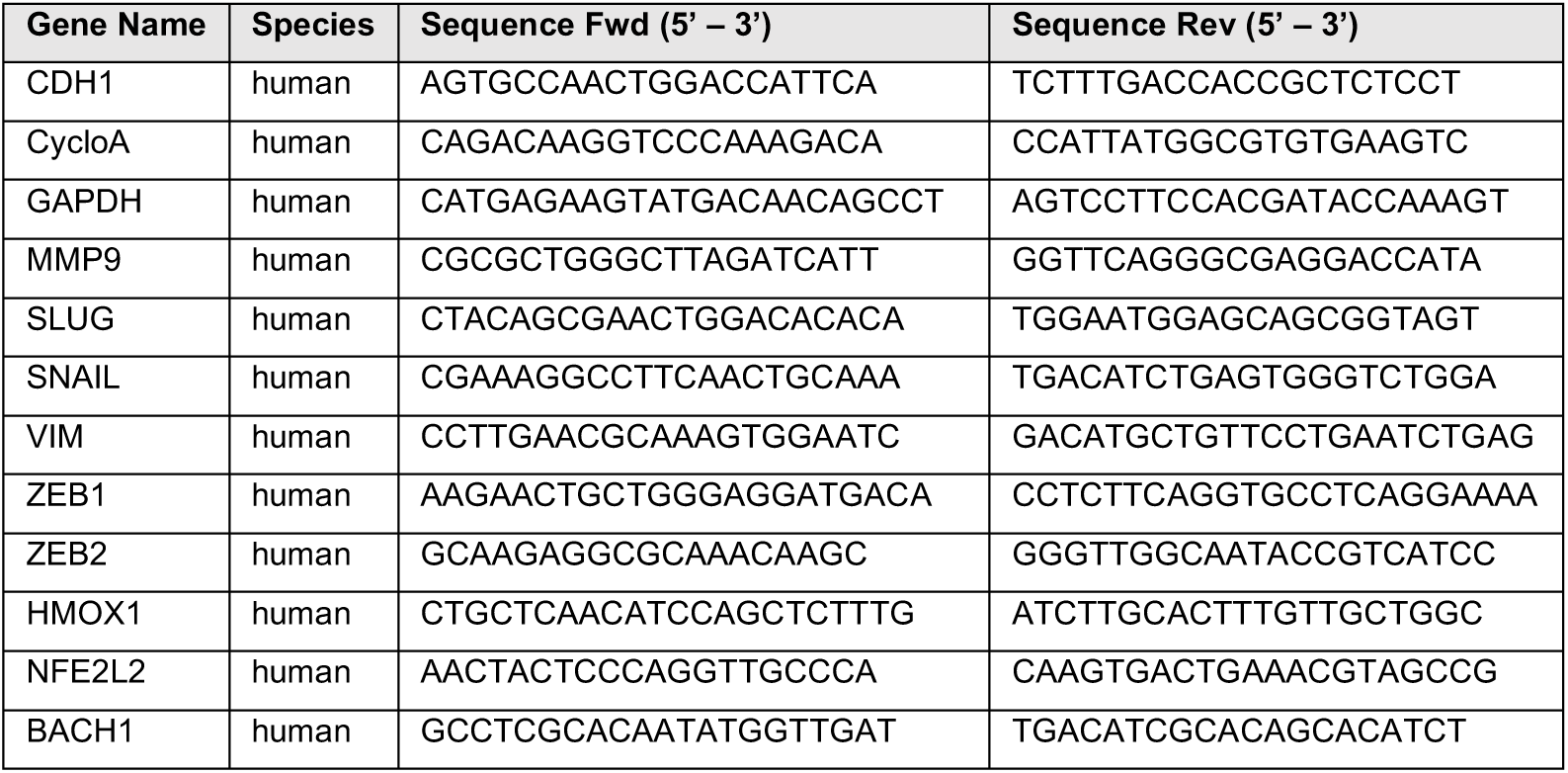
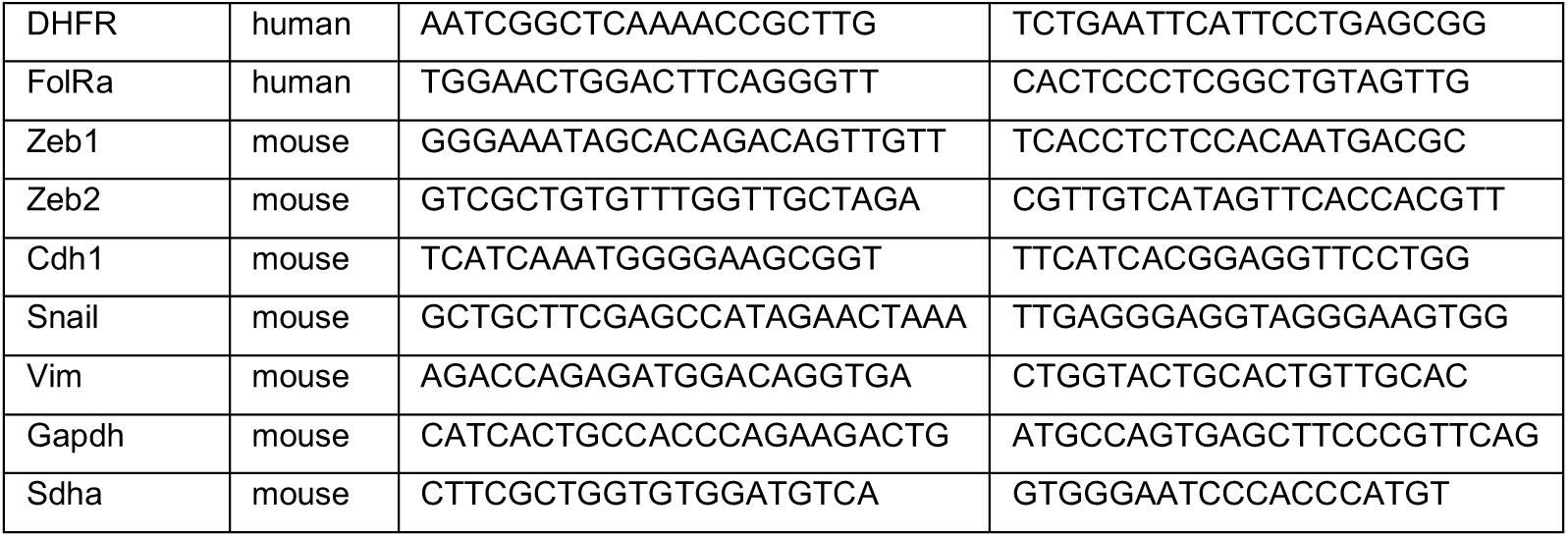

### shRNA-mediated Gene Silencing

Knockdown with shRNA was performed in MDA-MB-468 and 4T1 cells using lentiviral particles expressing two different shRNAs targeting the gene of interest and one non-silencing control vector (pGIPZ-shSCR: TTACTCTCGCCCAAGCGAG (Code: RHS4346). For silencing of the *MTHFD2* gene in MDA-MB-468 cells, pGIPZ shMTHFD2-2: ATTGCATTTCTATTGGCCT (V2LHS_90968) purchased from Horizon Discovery was used. For silencing of the *ALDH1L2* gene in MDA-MB-468 cells, pGIPZ shALDH1L2: ACTGCTTTATCAACATCCG (Code: V3LHS_385780) purchased from Horizon Discovery was used. For silencing of the *Mthfd1l* gene in 4T1 cells, pGIPZ shMTHFD1L-2: TAGATTTCAATTTCATCTG (Code: V2LHS_96542) purchased from Horizon Discovery was used. For silencing Mthfd2 gene in 4T1 cells, pGIPZ shMthfd2-2: ATGACAACGGCTTCATTTC (Code: V2LMM_23800) purchased from Horizon Discovery was used. Plasmids used for generation of lentivirus were amplified from glycerol stocks, isolated (Machery&Nagel plasmid isolation kit), and used for transfection of HEK293T cells. Lentivirus was produced for 24 h in HEK293T cells co-transfected with the viral core packaging construct pCMVR8.74, the VSV-G envelope protein vector pMD2.G and the respective pGIPZ-shRNA-target plasmid using lipofectamine 3000 (Invitrogen #L3000008). Filtered virus supernatant was used to transduce MDA-MB-468 and 4T1 cells in DMEM supplemented with 2% FBS. After 24 h incubation at 37°C, virus was removed and cells were cultured in DMEM supplemented with 10 % FBS for 48 h. 96 h post transduction, cells were cultured in selection media containing 2 μg/mL of puromycin (Sigma #P8833) for five passages to obtain stably transduced GFP-positive cells.

### Generation of KO cells

Gene knock-out (KO) using the CRISPR-Cas9 system was performed in MDA-MB-468 using a vector expressing hCas9 and the respective guideRNA to target the gene of interest. For silencing of the *MTHFD2* gene in MDA-MB-468 cells, two different gRNA sequences (gRNA#918: CGCCAACCAGGATCACACTC (Vector ID: VB190718-1104ncn); gRNA1.Ex: GCTCGCGGCAGTTCGGTAAGA (Vector ID: VB190719-1026vzh) purchased from VectorBuilder were used. A control vector was used scramble_gRNA (GTGTAGTTCGACCATTCGTG (Vector ID: VB190718-1103vja) together with both constructs. To generate a *MTHFD2*-KO cell line, 300.000 cells were seeded the day prior to transfection into a 6 well plate in DMEM supplemented with 10% FBS. 24 h later, the culture medium was replaced with 2 mL of DMEM-F12 (LONZA #12-719F) supplemented with 10 % FBS, Glucose (17 mM final concentration), Glutamine (2 mM final concentration), and 1 mM Sodium Formate and the cells were transfected using Opti-MEM, Lipofectamine, P3000 and 2 μg of plasmid DNA. Post incubation for 24 h, the transfection mix was removed and cells were replaced with fresh DMEM-F12. 72 h post transfection, cells were sorted into single cells by FACS based on their GFP fluorescence. Single cells were grown in DMEM-F12 supplemented with 1% Pen-Strep (Sigma, # P4333) until the formation of single colonies. These were further expanded and characterised by means of Western Blot.

### siRNA-mediated Gene Silencing

500.000 MDA-MB-468 cells were transfected with 40 pmol siRNA upon seeding in 6 well-plates using Lipofectamine 3000 (ThermoFisher) according to manufacturer’s instructions. Non-silencing control siRNA (siRNA ON-TARGETplus Human non-targeting (1810-10-05) siRNA SMARTpool) and siRNA directed against human ALDH1L2 mRNA (siRNA ON-TARGETplus Human ALDH1L2 (160428; 26918-01-0005) siRNA SMARTpool) were purchased from Dharmacon.

### Animal Model

Animal experiments were performed according to all applicable laws and regulations, after receiving approval by the institution’s Animal Experimentation Ethics Committee at UL (AEEC) and the veterinarian service of the Ministry of Agriculture, Viniculture & Rural Development (TumorMetab LUPA 2020/01). They ensure that care and use of animals for research purposes was conducted according to the EU Directive 2010/63/EU, as well as the Grand-Ducal Regulation of January 11, 2013 on the protection of animals used for scientific purposes. These included the justification of the use of animals, their welfare and the incorporation of the principles of the 3R’s (Replacement, Reduction and Refinement). A biostatistician reviewed all animal protocols. Mice were housed in a specific pathogen free (SPF) facility at a relative humidity of 40–70%, at 22 °C, and in 12 h dark/light cycles. Syngeneic 4T1 mammary carcinoma cells were orthotopically implanted into the left mammary fat pads (1 injection/mouse) of immune competent female Balb/c mice according to established protocols [64]. Briefly, each injection contained 2,000 cells in a mixture of 25 μL PBS and 25 μL matrigel. 9 mice per group were injected with either 4T1 SCR or 4T1 *MTHFD1L* KD cells. Primary tumor growth was monitored between day 7 and day 35 of the experiment. Weight was monitored and no weight loss was observed. Experiment was terminated after 6 weeks and lung and liver were prepared for examination of metastatic outgrowth. No metastases were found in the liver. Macroscopic lung metastases were blindly counted under a microscope and microscopic lung metastases were visualized by H&E staining.

#### H&E staining

H&E staining was performed on 10 μm sections of paraffin embedded lung tissue. Staining was performed according to established protocols. Briefly, selected sections were dehydrated with MeOH and stained with Gill 2 hematoxylin. Sections were neutralized with successive washes of tap water, hard water (10g MgSO4 and 0.7g NaHCO3 per L), and distilled water. Subsequently, sections were stained with Eosin-solution and dehydrated through successive washes with 80%, 95%, 100% EtOH and Xylol prior to mounting. Images for quantification were acquired using BioTek Cytation 5 Cell Imaging Multimode Reader. Area of lung and metastasis were measured in ImageJ. Representative images for publication were acquired using Olympus IX83 microscope. Collapsed lungs were excluded from quantitative analysis.

### Proteomics

SILAC (Stable Isotope Labeling with Amino acids in Cell culture) strategy was used for proteomic analysis. In general, MDA-MB-468 cells were cultivated in DMEM-F12 SILAC medium supplemented either with Lys^0^ and Arg^0^ (light channel) or Lys^8^ and Arg^10^ (heavy channel). After 6 passages, labeling efficiency of heavy channel was confirmed using LC-MS/MS. Cells in light channel were treated with 50 nM MTX for 24h and 72h, while heavy channel was used as control. Cell pellets of biological duplicates were collected. Proteins were extracted in lysis buffer (50mM ammonium bicarbonate, 6M Urea, 2M Thio-urea, pH 8) following a 30 min incubation at 4 °C in the presence of protease inhibitors (cOmplete™ EDTA-free Protease Inhibitor Cocktail, Roche). Following centrifugation at 16,000 g for 10 min, supernatants were taken for protein quantification. Samples from light channel (50 μg protein) were mixed with control heavy channel (50 μg protein) for protein reduction (5 mM DTT, 1 h incubation at 37°C) and alkylation (10 mM IAA, 45min in dark at room temperature). Protein digestion was performed with Lys-C (FUJIFILM Wako, 125-05061) at 1:30 ratio (enzyme/protein substances) for 4 h at 37 °C, then samples were diluted 4 times with 50 mM ammonium bicarbonate and digested overnight with 1 μg of trypsin at 37 °C. Digestion was terminated through addition of formic acid (1% final concentration). Digested peptides were cleaned up with reverse phase Sep-Pak C18 1 cc Vac Cartridge (Waters, WAT054955) and eluted with 1 mL 50% ACN. Eluted peptides were dried by Speedvac (Thermo Fisher Scientific) and re-suspended in 0.1% formic acid. Peptide concentration was measured with Nanodrop. Peptides were measured by LC-MS/MS on Q-Exactive HF mass spectrometer (Thermo Fisher) connected to a Dionex Ultimate 3000 (Thermo Fisher). 500 ng of peptides were loaded onto a trap column (Acclaim PepMap 75 μm × 2 cm, C18, 3 μm) and separated on a 25 cm Acclaim pepmap RSLC column (75 μm × 25 cm, C18, 2 μm) using a 150 min gradient (2% to 90% acetonitrile) with a flow rate of 0.3 μL/min. MS data were acquired in data dependent mode. Survey scans of peptide precursors from 375 to 1500 m/z were performed at 70,000 resolution with a 3×10^6^ ion count target and the top 12 abundant peaks from survey scan were selected for fragmentation. Tandem MS was performed by isolation at 1.4 m/z with the quadrupole, HCD fragmentation with a normalized collision energy of 28. The MS2 ion count target was set to 1×10^5^ and the max injection time was 45 ms. Only precursors with a charge state of 2–7 were sampled for MS2. The dynamic exclusion duration was set to 20 s with a 10 ppm mass tolerance around the selected precursor and its isotopes. Each sample was analyzed twice as technical replicates. All raw data was analyzed with MaxQuant (version 1.6.7.0) and searched with Andromeda against the Homo sapiens database from Uniprot. The minimal peptide length was set to 7 amino acids and the maximum of 3 missed cleavages were allowed. The search included variable modifications of methionine oxidation and N-terminal acetylation, deamidation (N and Q) and fixed modification of carbamidomethyl cysteine. The “Match between run” was checked within 1 min retention time window. Mass tolerance for peptide precursor and fragments were set as 10 ppm and 20 ppm, respectively. The FDR was set to 0.01 for peptide and protein identifications. SILAC based protein quantification (MaxQuant built-in) was used for quantitative evaluation of identified proteins. ProTIGY (https://github.com/broadinstitute/protigy), an R based tool, was used for differential analysis of MaxQuant output.

### Statistics

Unpaired *t*-test with Welch’s correction was applied for pairwise comparison (two-sided) using GraphPad Software Vers.8. For comparison of multiple groups ordinary Brown Forsythe & Welch one-way or two-way ANOVA with Dunnett’s multiple comparison or Games-Howell’s multiple comparisons test was performed using GraphPad Software Vers.8. Resulting p-values for comparisons of interest are given as numerical values within the respective graphs. For normalization, data points of one experiment were either normalized to the untreated control or divided by the global mean of the individual experiment. We define one *n* as one independent biological experiment (in some cases further consisting of several wells, e.g. triplicate wells for all stable isotope tracing experiments). The technical mean of one biological experiment was considered as one *n*. The mean values of several independent, biological experiments (as indicated in figure legends) were plotted and used for statistical analysis as indicated.

## Data availability

The data that support the findings of this study are available from the corresponding author upon reasonable request. Proteomics data (Fig S2F) are deposited on a server with the following pride ID: PXD027175. At the current moment the access is set to private. It will be made public after acceptance of the manuscript. Access credentials for reviewers are as follows:

**Password:** 5Mhn9v58

