## Supplemental Figures Kiweler et al_012022 for "Mitochondria preserve an autarkic one-carbon cycle to confer growth-independent cancer cell migration and metastasis"

### Supplementary Figure 1

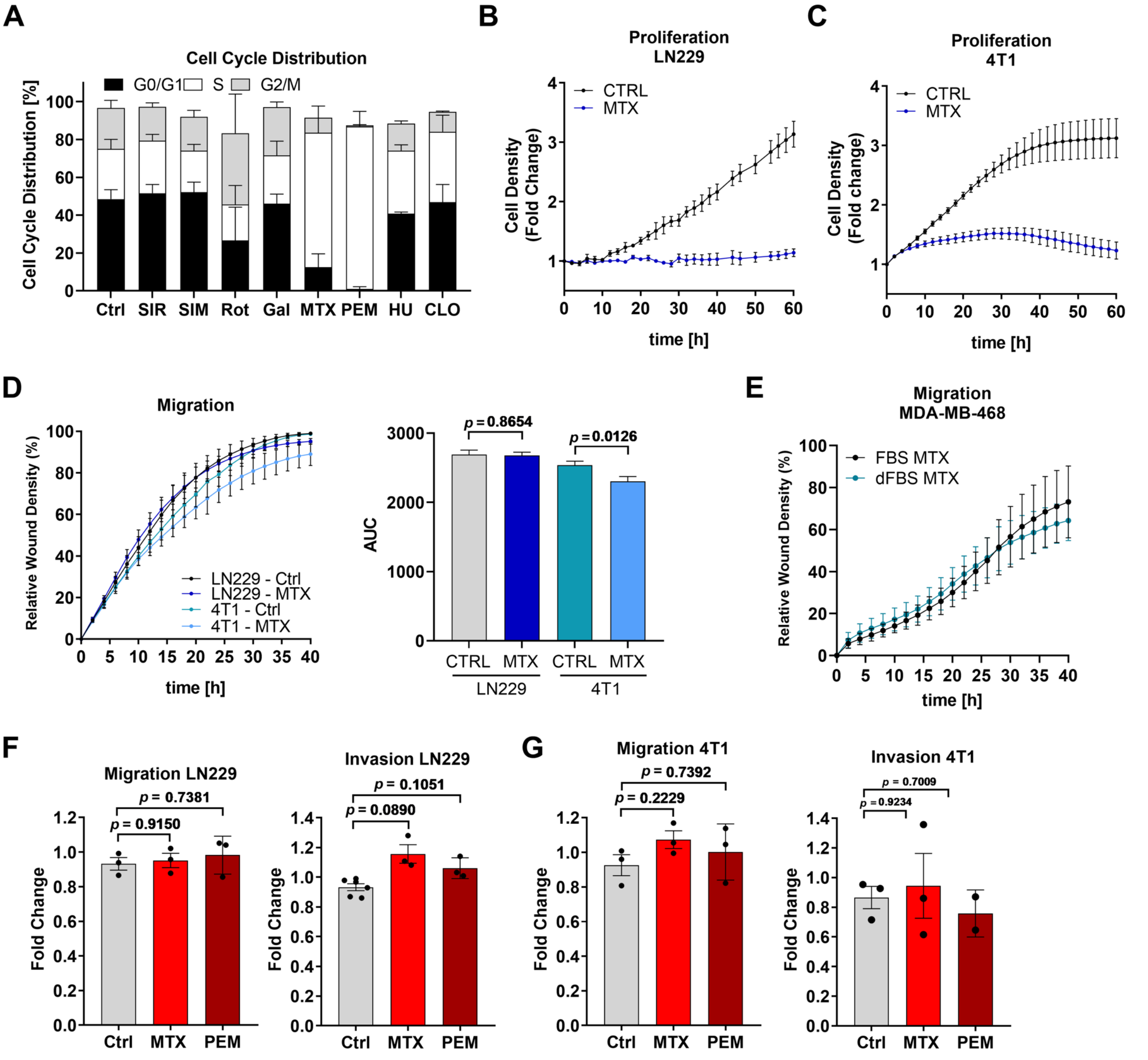


**Supplementary Figure 1: (A)** MDA-MB-468 cells were treated for 48 h with 100 nM Sirolimus (SIR), 1 µM Simvastatin (SIM), 50 nM Rotenone (Rot), galactose (Gal) supplementation, 50 nM Methotrexate (MTX), 1 µM Pemetrexed (PEM), 0.5 mM hydroxyurea (HU), and 100 nM Clofarabine (CLO). Cells were analyzed for cell cycle distribution by flow cytometry of PI-stained cells; mean ± SD (n = 3 - 11). **(B)** Proliferation of LN229 cells in response to 50 nM MTX; mean ± SEM (n = 3). **(C)** Proliferation of 4T1 cells in response to 75 nM MTX; mean ± SEM (n = 4). **(D)** Migration of LN229 and 4T1 cells in response to 50 (LN229) or 75 (4T1) nM MTX and respective AUC; mean ± SEM (n = 5); unpaired t-test with Welch's correction. (**E**) Migration of MDA-MB-468 cells after 24 h treatment with 50 nM MTX in medium supplemented with normal or dialyzed FBS; graph shows mean ± SEM of 2 independent experiments. **(F), (G)** Migration and invasion of LN229 (F) and 4T1 (G) cells after 24h treatment with 50 nM (F), 75 nM (G) MTX was assessed using ECM-Collagen-coated or non-coated Boyden chambers. Each dot represents an independent experiment; mean ± SEM; Brown-Forsythe and Welch one-way ANOVA with Dunnett's multiple comparisons test.

#### Supplementary Figure 2

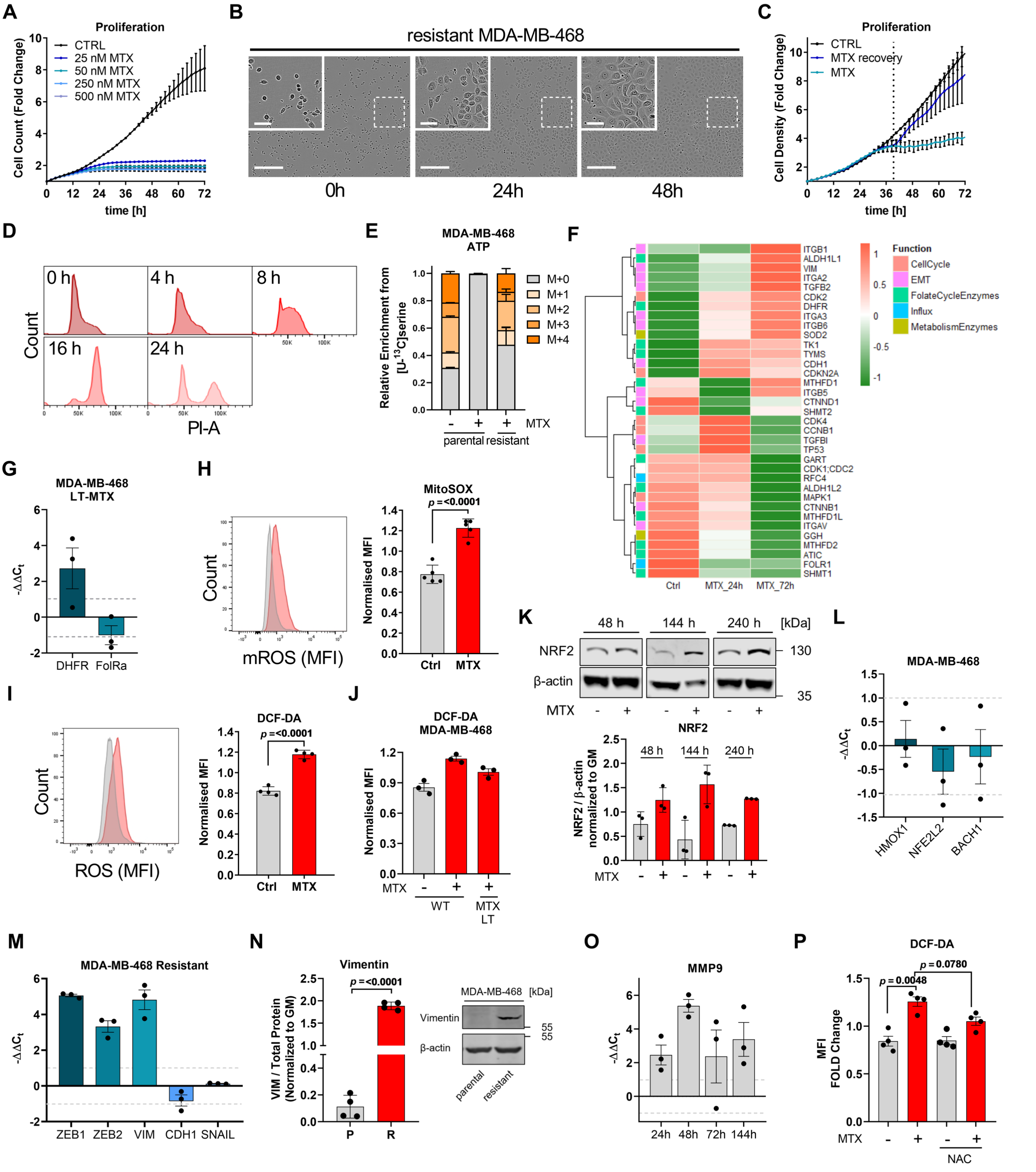


**Supplementary Figure 2: (A)** Proliferation of NucLight Rapid Red stained MDA-MB-468 cells upon the indicated concentrations of MTX; mean ± SEM (n = 3). **(B)** Morphology of proliferating MTX-resistant MDA-MB-468 cells. Bright-field images are representative of independent experiments. Scale bars correspond to 60 and 300 µm. **(C)** Growth of MDA-MB-468 cells in response to 50 nM MTX and growth recovery upon removal of MTX after 38 h assessed as fold change of cell density; mean ± SEM (n = 3). **(D)** Recovery of cell cycle profile at the indicated time points after MTX removal as in (C). Histograms are representative of three independent experiments. **(E)** MID of intracellular ATP upon [U-$^{13}$C]serine tracer in response to 24 h 50 nM MTX in parental MDA-MB-468 cells and MTX-resistant MDA-MB-468 cells. Graph shows mean ± SEM of two independent experiments each measured in triplicate wells. **(F)** Heatmap of selected protein expressions from SILAC quantitative proteomics measurement with different MTX treatment durations. The log2 fold changes of each protein were standardized by row. Molecular function of proteins are indicated with colors. **(G)** mRNA expression from the indicated genes was quantified by real-time RT-qPCR in resistant long-term (LT) MTX treated MDA-MB-468 cells relative to MDA-MB-468 WT cells. (**H**) Mitochondrial ROS levels in response to 24 h 50 nM MTX measured by flow cytometric quantification of MitoSOX mean fluorescence intensity. Each dot represents an independent experiment; mean ± SD; unpaired t-test with Welch's correction. **(I)** Intracellular ROS levels in response to 24 h 50 nM MTX measured by flow cytometric quantification of DCF-DA mean fluorescence intensity. Each dot represents an independent experiment; mean ± SD; unpaired t-test with Welch's correction. **(J)** Intracellular ROS levels in response to 24 h MTX treatment in MDA-MB-468 WT cells and in LT MTX treated MTX_resistant MDA-MB-468 cells measured by flow cytometric quantification of DCF-DA mean fluorescence intensity. Each dot represents one independent experiment; mean ± SD. (**K**) NRF2 expression in MDA-MB-468 cells after treatment with 50 nM MTX for the indicated time was quantified by Western Blot. Normalized signal intensity was quantified relative to β-actin signal. Each dot represents an independent experiment; mean ± SD. (**L**) mRNA expression from the indicated target genes in MDA-MB-468 cells was measured after 144 h 50 nM MTX and quantified relative to untreated cells using real-time RT-qPCR. Each dot represents one independent experiment; mean ± SD. (**M**) mRNA expression from the indicated target genes in MTX-resistant MDA-MB-468 cells relative to parental MDA-MB-468 cells as measured using real-time RT-qPCR. Each dot represents an independent experiment; mean ± SEM. **(N)** Expression of vimentin in parental and MTX-resistant MDA-MB-468 cells; β-actin serves as loading control. Quantification of vimentin signal intensity relative to total protein stain. Each dot represents an independent experiment; mean ± SD; unpaired t-test with Welch's correction. (**O**) mRNA expression of MMP9 was quantified in MDA-MB-468 cells after 50 nM MTX at the indicated time points using real-time RT-qPCR. Each dot represents an independent experiment; mean ± SEM. **(P)** Intracellular ROS levels in MDA-MB-468 cells in response to 24 h 50 nM MTX and 10 mM NAC as measured by flow cytometric quantification of DCF-DA mean fluorescence intensity. Each dot represents an independent experiment. Graph shows mean ± SD; Brown-Forsythe and Welch ANOVA test with Games-Howell's multiple comparisons test.

### Supplementary Figure 3

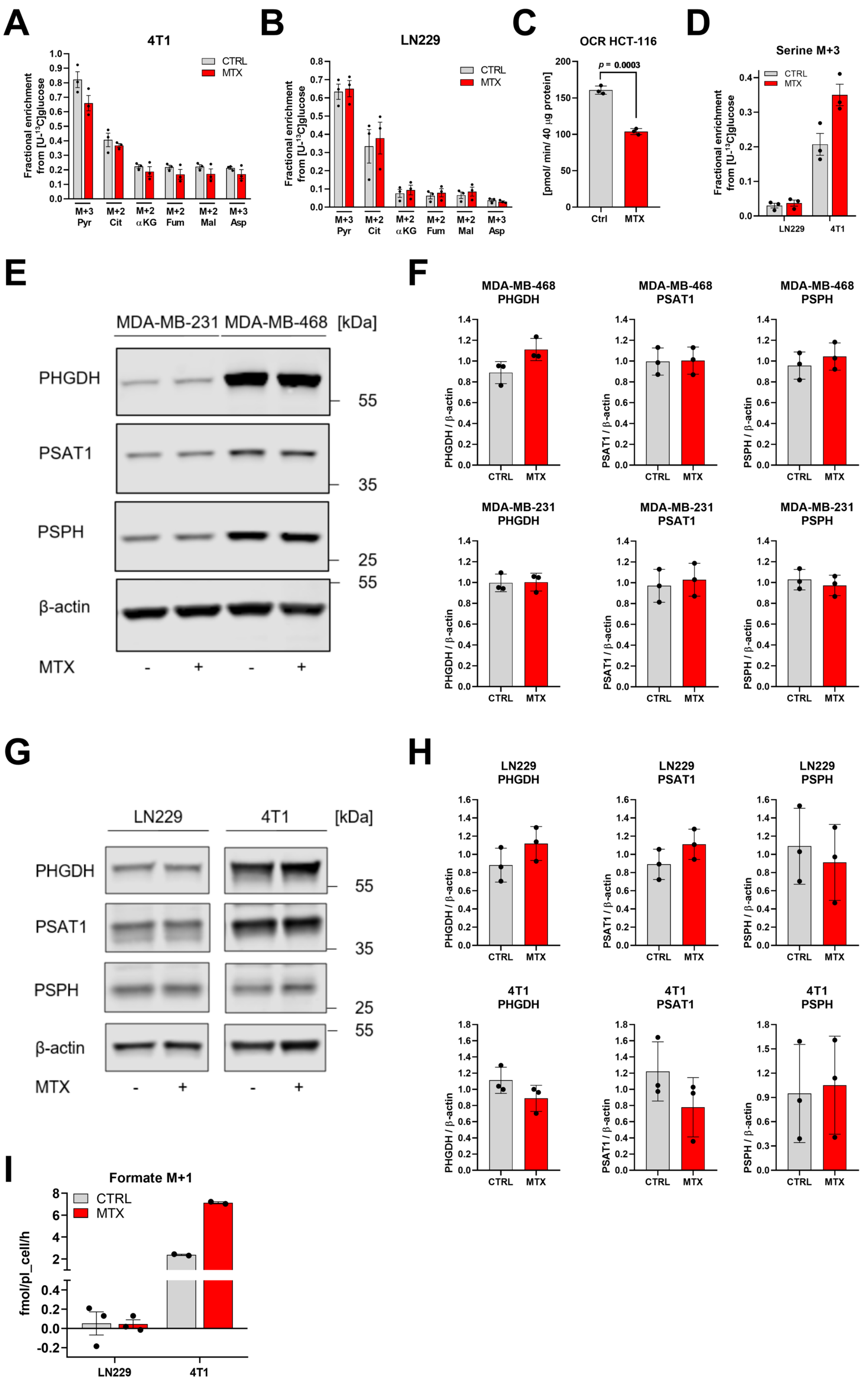

**Supplementary Figure 3: (A), (B)** Relative isotopologue abundance following [U-$^{13}$C]glucose tracing in 4T1 (A) and LN229 (B) cells in response to 24 h 75 nM MTX (A) and 50 nM MTX (B). Each dot represents an individual experiment composed of triplicate wells; mean ± SEM. **(C)** Basal cellular respiration in response to 24 h 50 nM MTX treatment was determined in HCT-116 cells as quantification of mitochondrial OCR. Each dot represents an individual experiment composed of six technical replicates; mean ± SEM; unpaired t-test with Welch's correction. **(D)** M+3 isotopologue abundance of serine upon [U-$^{13}$C]glucose tracing in response to 24 h 50 nM MTX in LN229 and 24 h 75 nM MTX in 4T1 cells. Each dot represents an independent experiment in triplicate wells; mean ± SEM. **(E)** Expression of PHGDH, PSAT1, and PSPH in MDA-MB231 and MDA-MB-468 cells upon 24 h 50 nM MTX treatment; β-actin serves as loading control. **(F)** Treatment as in (E). Quantification of PHGDH, PSAT1, and PSPH Western blot signal intensity relative to β-actin normalized to global mean in MDA-MB231 and MDA-MB-468 cells. Each dot represents an independent experiment; mean ± SD. **(G)** Expression of PHGDH, PSAT1, and PSPH in LN229 and 4T1 cells upon 24 h 50 nM MTX (LN229) and 75 nM MTX (4T1) treatment; β-actin serves as loading control. **(H)** Treatment as in (G). Quantification of PHGDH, PSAT1, and PSPH Western blot signal intensity relative to β-actin and normalized to global mean in LN229 and 4T1 cells. Each dot represents an independent experiment; mean ± SD. **(I)** M+1 formate release rate in LN229 and 4T1 cells after 24 h [U-$^{13}$C]glucose tracing and treatment with 50 nM (LN229) and 75 nM (4T1) MTX. Each dot indicates an independent experiment measured in triplicate wells; mean ± SEM.

#### Supplementary Figure 4

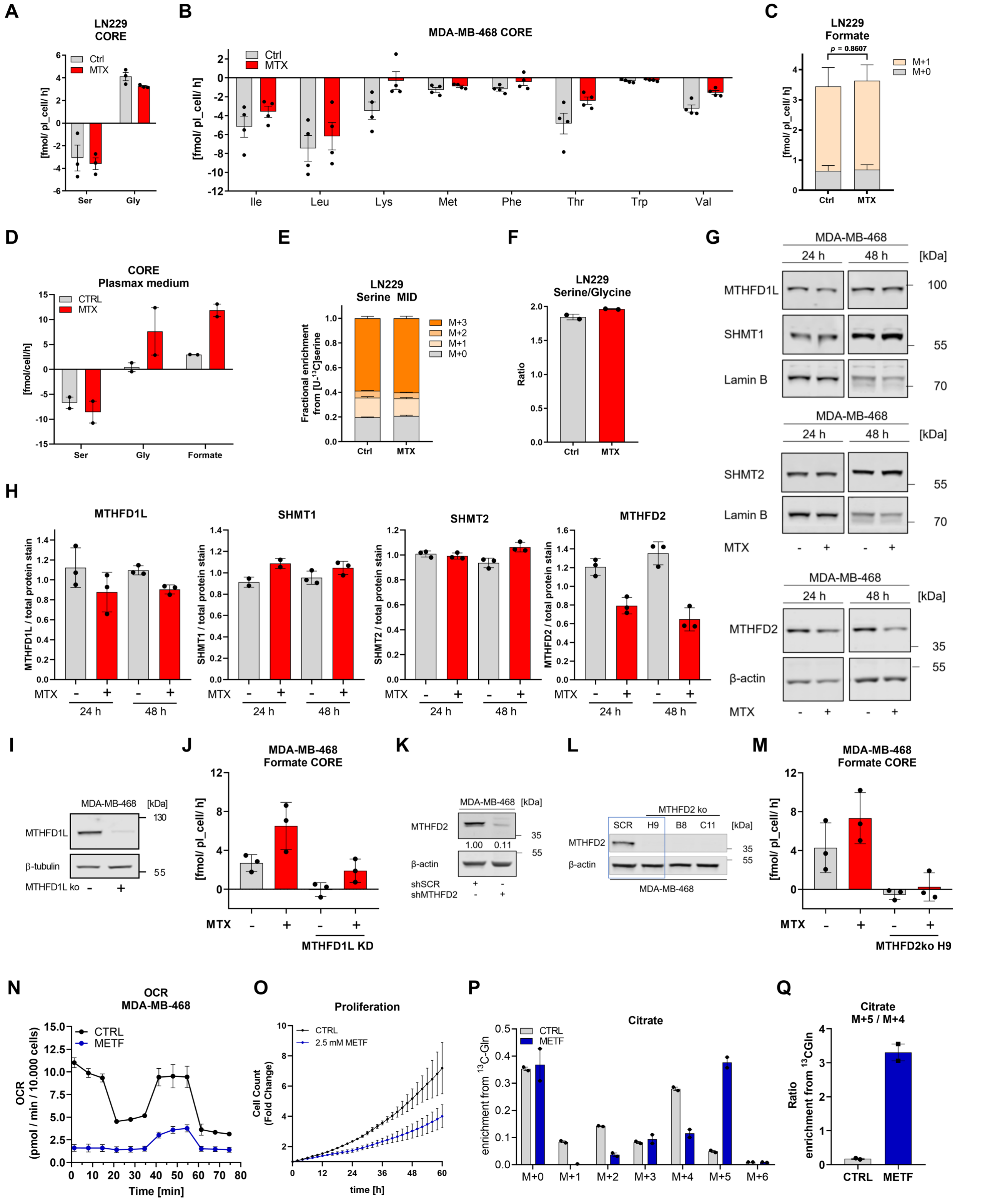

**Supplementary Figure 4: (A)** Absolute CORE rates of serine and glycine of LN229 cells in response to 24 h 50 nM MTX. Each dot represents the mean of an individual experiment each measured in triplicate wells; mean ± SEM. **(B)** Absolute CORE rates of essential amino acids as indicated in response to 24 h 50 nM MTX. Each dot represents an independent experiment in triplicate wells; mean ± SEM. **(C)** Formate release rates of M+0 and M+1 formate isotopologues upon [U-$^{13}$C]serine tracing in LN229 cells in response to 24 h 50 nM MTX. Graph shows mean ± SEM of three independent experiments each measured in triplicate wells; unpaired t-test with Welch's correction. **(D)** Absolute CORE rates of serine, glycine, and formate in MDA-MB-468 cells in response to 24 h 50 nM MTX in Plasmax medium. Each dot represents an independent experiment in triplicate wells; mean ± SEM. **(E)** MID of intracellular serine upon [U-$^{13}$C]serine tracing in LN229 cells in response to 24 h 50 nM MTX; mean ± SEM of two independent experiments each measured in triplicate wells. **(F)** Ratio of intracellular serine and glycine levels in response to 24 h 50 nM MTX in LN229 cells. Each dot represents an independent experiment in triplicate wells; mean ± SEM. **(G)** Expression of MTHFD1L, SHMT1, SHMT2, and MTHFD2 in MDA-MB-468 cells 24 h and 48 h 50 nM MTX; β-actin serves as loading control. **(H)** Treatment as in (G). Quantification of MTHFD1L, SHMT1, SHMT2, and MTHFD2 Western blot signal intensity relative to total protein stain normalized to GM in MDA-MB-468 cells. Each dot represents an independent experiment; mean ± SD. **(I)** Residual MTHFD1L protein expression in MDA-MB-468 MTHFD1L CRISPR KO cells was determined on Western Blot. β-tubulin serves as loading control. **(J)** Absolute CORE rates of formate from mock or shMTHFD1L-transfected MDA-MB-468 cells upon 24 h 50 nM MTX. Dots represent technical replicates of a representative experiment; mean ± SD. **(K)** Residual MTHFD2 protein expression in mock or shMTHFD2-transfected MDA-MB-468 cells was determined on Western Blot. β-actin serves as loading control. **(L)** Residual MTHFD2 protein expression in three MDA-MB-468 MTHFD2 CRISPR KO clones was determined on Western Blot. β-actin serves as loading control. **(M)** Absolute CORE rates of formate from MDA-MB-468 cells upon depletion of MTHFD2 (CRISPR clone H9) and 24 h 50 nM MTX. Dots represent technical replicates of a representative experiment; mean ± SD. **(N)** Oxygen consumption rate (OCR) in MDA-MB-468 cells after 24h treatment with 2.5 mM Metformin. Graph shows mean ± SD of one representative experiment measured in six technical replicates. **(O)** Proliferation of MDA-MB-468 cells in response to treatment with 2.5 mM Metformin. Graph shows mean ± SEM of two independent experiment. **(P)** Citrate MID from [U-$^{13}$C]Glutamine tracing in MDA-MB-468 cells after 24 h treatment with 2.5 mM Metformin. Graph shows mean ± SD of two independent experiments. **(Q)** Ratio of M+5/M+4 istotopologues of citrate after 24h treatment with 2.5 mM Metformin and tracing with [U-$^{13}$C]Glutamine tracing. Graph shows mean ± SD of two independent experiments.

### Supplementary Figure 5

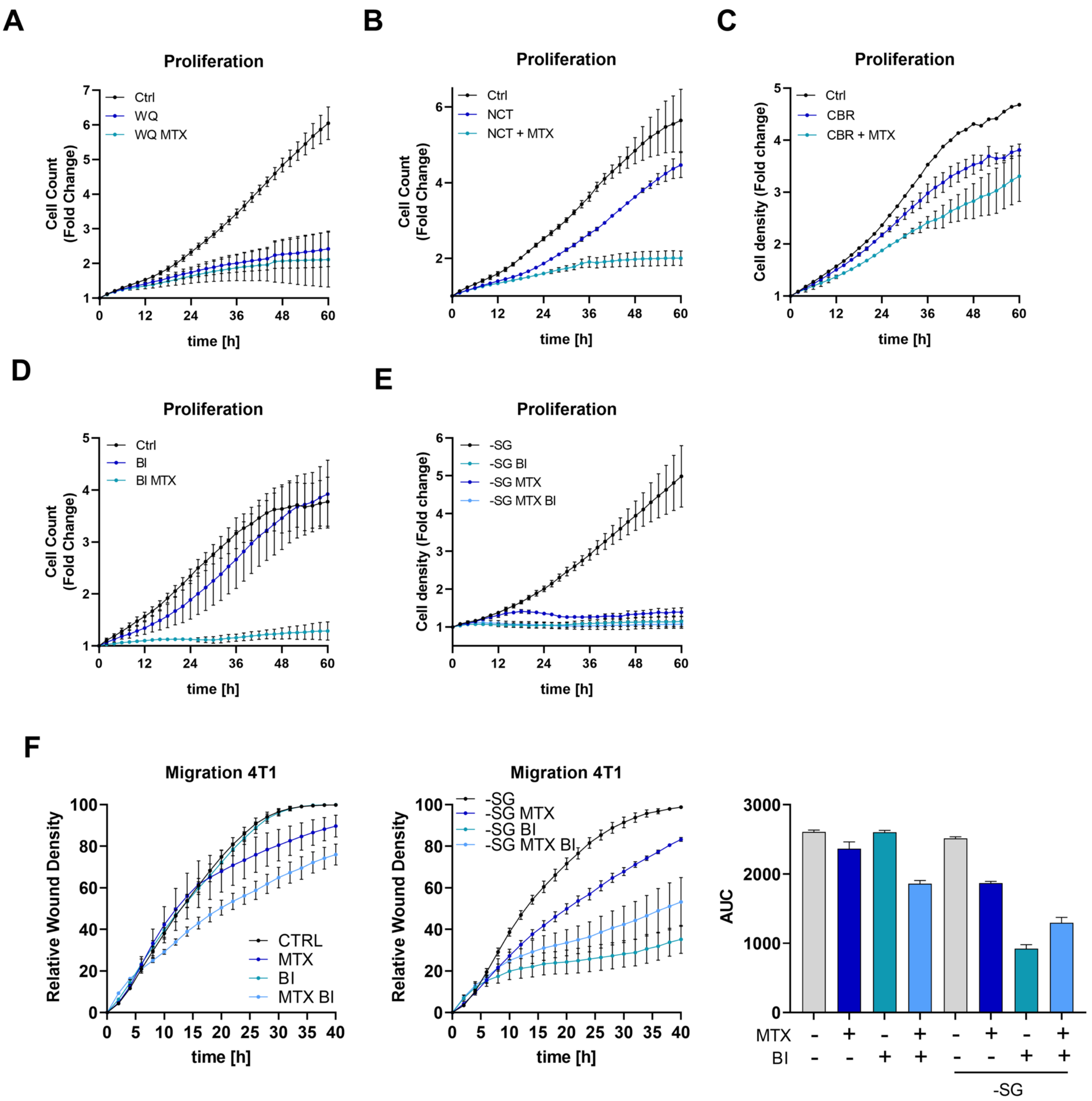


**Supplementary Figure 5: (A)** Proliferation of NucLight Rapid Red stained MDA-MB-468 cells upon treatment with 10 µM WQ-2101 and 50 nM MTX as indicated; mean ± SEM (n = 6 - 7). **(B)** Proliferation of NucLight Rapid Red stained MDA-MB-468 cells upon treatment with 10 µM NCT-502 and 50 nM MTX as indicated; mean ± SEM (n = 4). **(C)** Proliferation of MDA-MB-468 cells measured as fold cell density upon treatment with 30 µM CBR-5884 and 50 nM MTX as indicated; mean ± SEM (n = 1 - 2). **(D)** Proliferation of NucLight Rapid Red stained MDA-MB-468 cells upon treatment with 15 µM BI-4916 and 50 nM MTX as indicated; mean ± SEM (n = 2 - 3). **(E)** Proliferation of serine- and glycine-starved MDA-MB-468 cells measured as fold cell density upon treatment with 15 µM BI-4916 and 50 nM MTX as indicated; mean ± SEM (n = 3). (**F**) Migration of 4T1 cells upon treatment with 75 nM MTX and 15 µM BI in the presence or absence of serine and glycine in culture medium and respective AUC over 40 h. Graph shows mean ± SD of one representative experiment performed in 8 technical replicates (three independent experiments in total).

### Supplementary Figure 6

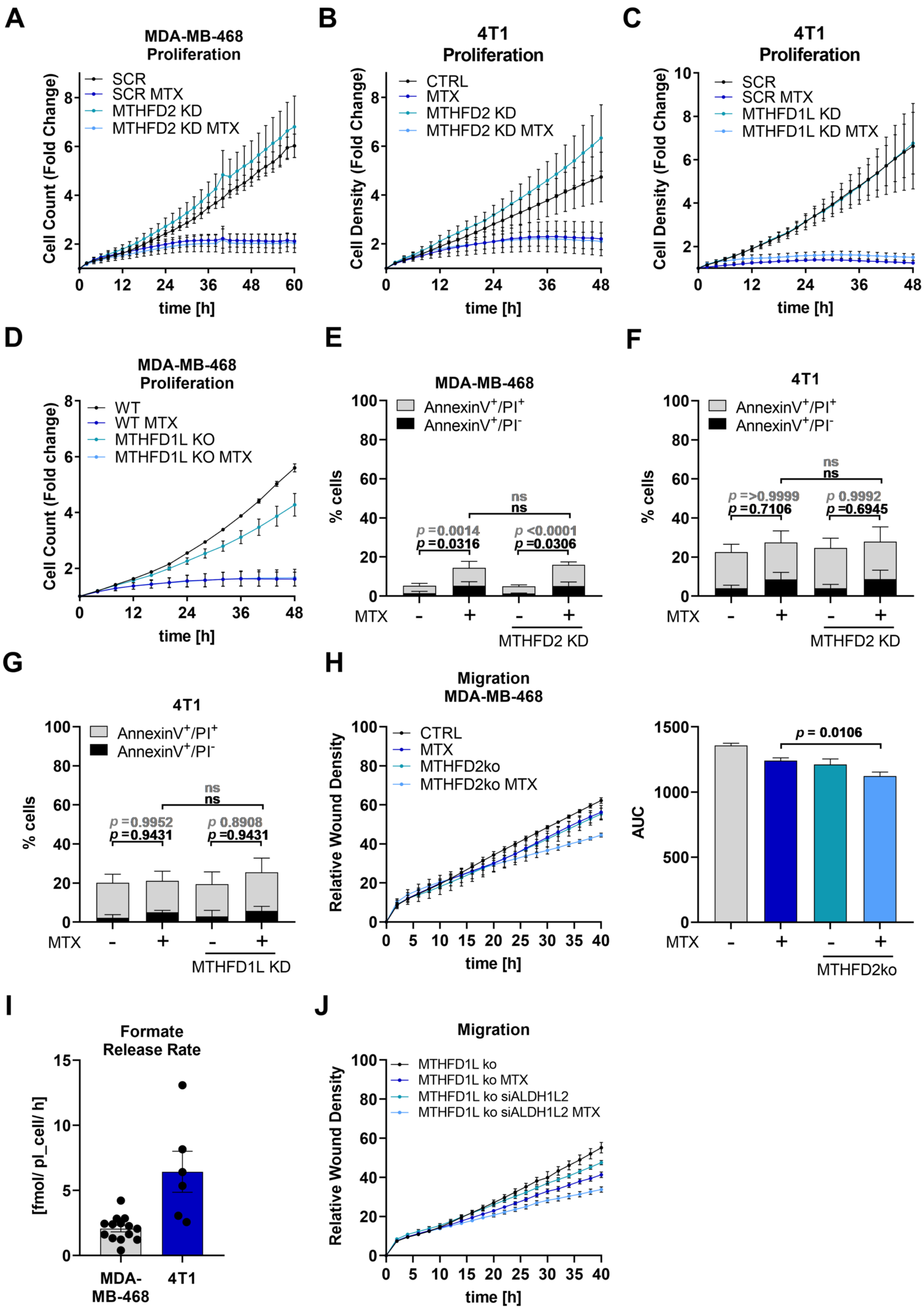


**Supplementary Figure 6: (A-D)** Proliferation of MDA-MB-468 (A,D) and 4T1 (B,C) cells upon MTHFD1L and MTHFD2 depletion as indicated after treatment with 50 nM MTX (A,D) or 75 nM MTX (B,C). Graph shows mean ± SEM (n = 6 (A),

1-4 (B), 3 (C), 4 (D)). (**E-G**) MDA-MB-468 (E) and 4T1 (F,G) cells depleted for MTHFD2 or MTHFD1L were treated with 50 nM (E) or 75 nM (F, G) MTX for 48h. Cell death was assessed by flow cytometry and AnnexinV-FITC/PI-staining; mean ± SD (n = 4). 2-way ANOVA with Dunnett's multiple comparisons test. **(H)** Migration of MDA-MB-468 cells (clone H9) in response to MTHFD2 knockout and 50 nM MTX treatment. Graph shows mean ± SEM (n = 3); Brown-Forsythe and Welch one-way ANOVA with Dunnett's multiple comparisons test. **(I)** Absolute formate release from MDA-MB-468 and 4T1 cells. Graph shows mean ±SEM. Each dot indicates an independent experiment performed in technical triplicates. **(J)** Migration of MDA-MB-468 MTHFD1L knockout cells in response to siRNA-mediated knockdown of ALDH1L2 and 50 nM MTX treatment (or respective shscramble control). Graph shows mean ± SEM of technical replicates (n = 7–8 wells).

### Supplementary Figure 7

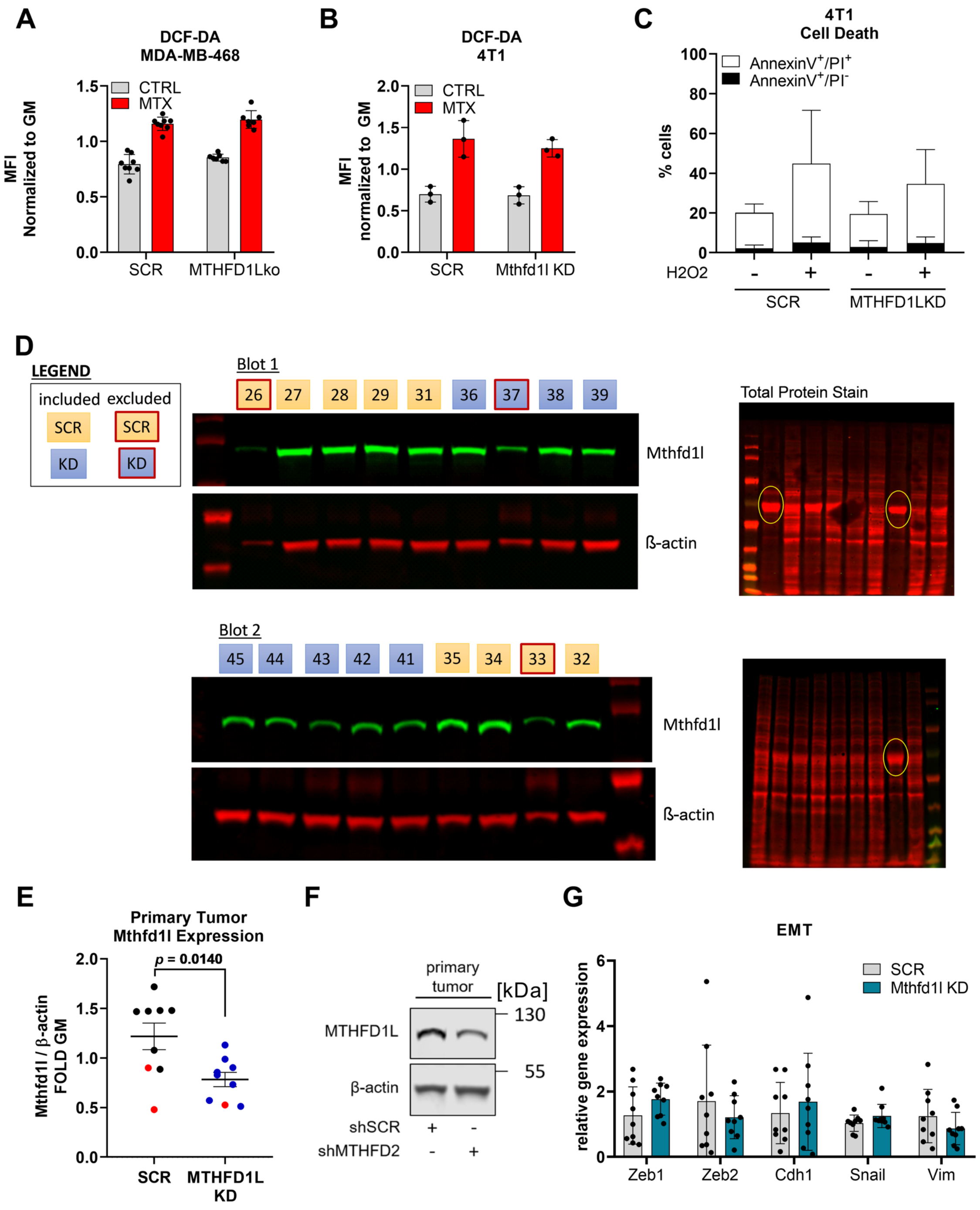


**Supplementary Figure 7: (A, B)** Intracellular ROS levels in MDA-MB-468 and 4T1 cells in response to 24 h 50 nM MTX (A) or 75 nM MTX (B) measured by flow cytometric quantification of DCF-DA mean fluorescence intensity. Each dot represents one independent experiment; mean ± SD. **(C)** 4T1 cells depleted or not for Mthfd1l were treated with 500 µM $H_2O_2$ for 24 h. Cell death was assessed by flow cytometry and AnnexinV-FITC/PI-staining; mean ± SD (n = 4). **(D)** Western blot images of protein lysates from primary tumor tissue. Indicated replicates were excluded from analysis in Figure 7D, due to the here presented finding that total protein in these samples was mainly composed of a single protein (see Total Stain). This indicates unreliability of the sample. (E) Mthfd1l expression in primary tumor tissue obtained by orthotopic implantation of 4T1 SCR and Mthfd1lKD breast cancer cells. Red dots indicate data points that were excluded as explained in (E). (**G)** Representative Western Blot image of Mthfd1l protein expression in primary tumor tissue in mice after orthotopic implantation of 4T1 scramble or Mthfd1l shRNA transfected breast cancer cells. β-actin serves as loading control. (**D**) mRNA expression from indicated target genes in primary tumor tissue in mice after orthotopic implantation of 4T1 scramble or Mthfd1l shRNA transfected breast cancer cells measured by real-time RT qPCR. Each dot indicates one individual animal; mean ± SD.
